# Alpha Synuclein Modulates Mitochondrial Ca^2+^ Uptake from ER During Cell Stimulation and Under Stress Conditions

**DOI:** 10.1101/2023.04.23.537965

**Authors:** Meraj Ramezani, Alice Wagenknecht-Wiesner, Tong Wang, David A. Holowka, David Eliezer, Barbara A. Baird

**Affiliations:** Department of Chemistry and Chemical Biology, Cornell University, Ithaca, New York 14853; Department of Biochemistry, Weill Cornell Medicine, New York, NY 10065

## Abstract

Alpha synuclein (a-syn) is an intrinsically disordered protein prevalent in neurons, and aggregated forms are associated with synucleinopathies including Parkinson’ disease (PD). Despite the biomedical importance and extensive studies, the physiological role of a-syn and its participation in etiology of PD remain uncertain. We showed previously in model RBL cells that a-syn colocalizes with mitochondrial membranes, depending on formation of N-terminal helices and increasing with mitochondrial stress.^1^ We have now characterized this colocalization and functional correlates in RBL, HEK293, and N2a cells. We find that expression of a-syn enhances stimulated mitochondrial uptake of Ca^2+^ from the ER, depending on formation of its N-terminal helices but not on its disordered C-terminal tail. Our results are consistent with a-syn acting as a tether between mitochondria and ER, and we show increased contacts between these two organelles using structured illumination microscopy. We tested mitochondrial stress caused by toxins related to PD, 1-methyl-4-phenyl-1,2,3,6-tetrahydropyridine (MPTP/MPP+) and carbonyl cyanide m-chlorophenyl hydrazone (CCCP), and found that a-syn prevents recovery of stimulated mitochondrial Ca^2+^ uptake. The C-terminal tail, and not N-terminal helices, is involved in this inhibitory activity, which is abrogated when phosphorylation site serine-129 is mutated (S129A). Correspondingly, we find that MPTP/MPP+ and CCCP stress is accompanied by both phosphorylation (pS129) and aggregation of a-syn. Overall, our results indicate that a-syn can participate as a tethering protein to modulate Ca^2+^ flux between ER and mitochondria, with potential physiological significance. A-syn can also prevent cellular recovery from toxin-induced mitochondrial dysfunction, which may represent a pathological role of a-syn in the etiology of PD.

## INTRODUCTION

Parkinson’s disease (PD) is the second most common neurodegenerative disorder in humans, increasing markedly with age and characterized by formation of Lewy bodies (LBs) and Lewy neurites (LNs) in the dopaminergic neurons of the brain substantia nigra.^2^ Alpha-synuclein (a- syn), an abundant presynaptic protein,^3^ is found in a filamentous form in LBs and LNs and is in other ways genetically and pathologically linked to PD and other synucleinopathies.^4^ Polymeropoulos et al. first reported a PD-related G209A mutation in the SNCA gene encoding for a- syn.^5^ Other studies further linked the SNCA gene and expressed a-syn variants to PD.^6–9^ Earlier onset of PD and a more severe course have been observed in patients with duplication or triplication of SNCA.^10^

A-syn is a 140 residue protein, found predominantly in neurons and characterized as intrinsically disordered in solution.^3^ However, a-syn has been shown to adopt a highly helical structure in the presence of negatively charged lipid surfaces.^11–15^ An extended helix forms upon binding to negatively charged phospholipid vesicles in the N-terminal amphipathic region (residues 1–100), and a broken helix forms when binding to phospholipid micelles.^12, 16–19^ The broken helix form of a-syn comprises helix-1 (residues 3 to 38) and helix-2 (residues 46 to 93), loosely connected by an unstructured flexible motif known as the linker region (residues 39 to 45) (Figure 1).^20^ Georgieva et al. proposed that the broken helix form of a-syn can serve as a tether between two phospholipid membranes,^21^ such as between synaptic vesicles and plasma membrane or two organellar membranes, and others have adopted this model as well.^22^ The C-terminal segment of a- syn (residues 100–140) is acidic, glutamate-rich, and remains disordered even in the presence of membranes.^16, 23^ This segment has been implicated in several protein interactions^24–28^ and contains residues that are targets for post translational modifications, notably phosphorylation of serine residue 129 (pS129). Only a small fraction of a-syn (less than 4%) is phosphorylated in normal brain tissue, but a dramatic accumulation of pS129 (More than 90%) is observed within LBs.^29, 30^

**Figure 1.**
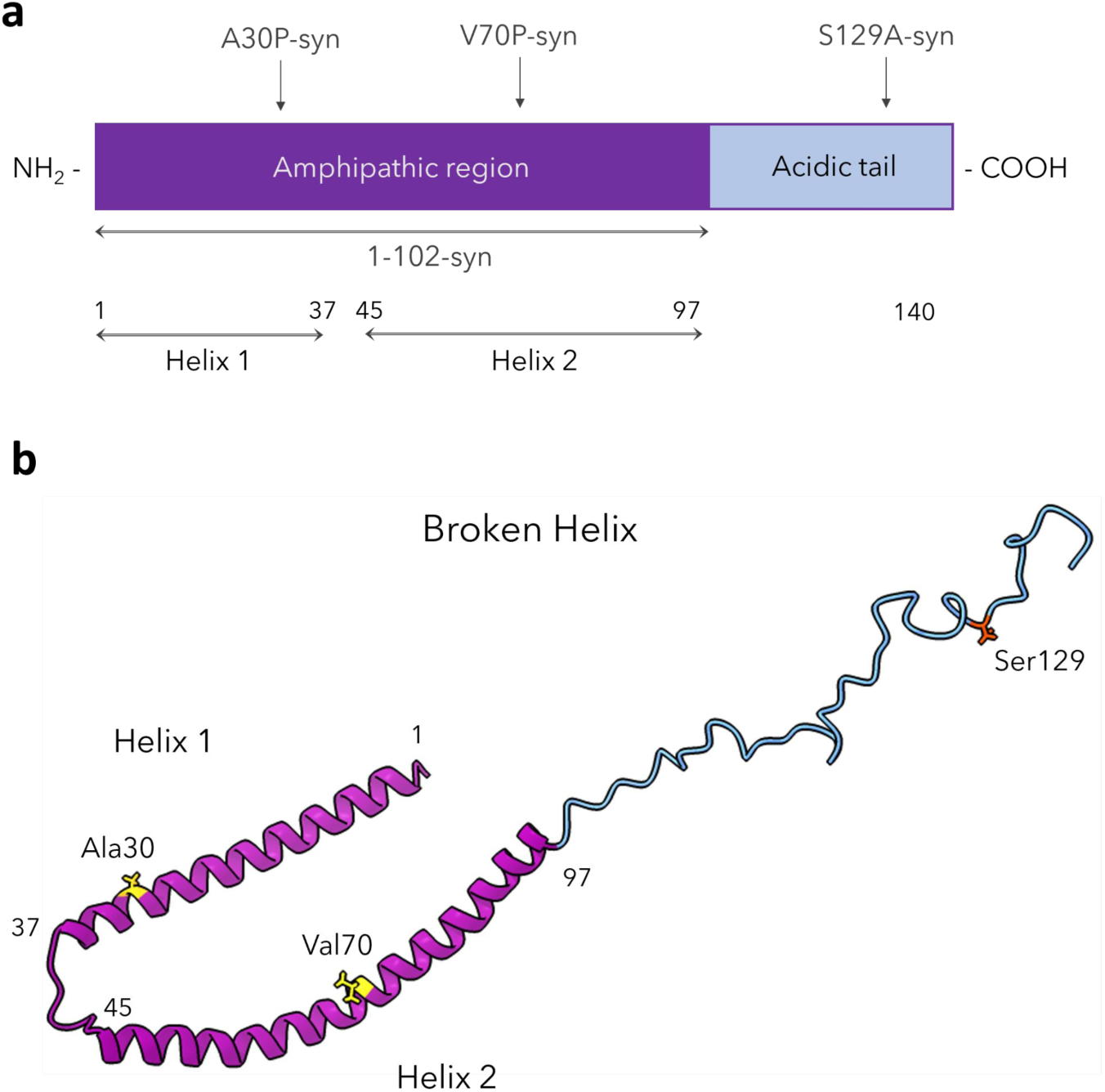
Structural features and variants of a-syn. **a)** Schematic representation of the a-syn primary sequence delineating the amphipathic membrane binding domain (purple) and the acidic C-terminal tail (blue); locations are indicated for helix-1 and helix-2 of the broken-helix state and for sites of mutations examined in the manuscript. **b)** Wt a-syn in the broken helix conformation (RCSB protein data bank entry 1XQ8) with locations shown for sidechains Ala 30, Val 70, and Ser 129, which were mutated in this study.

We previously established that RBL cells expressing human a-syn variants serve as a versatile model for evaluating intracellular distributions of a-syn and accompanying effects on cell function that are mediated by its membrane interactions.^1, 31^ RBL cells have internal structures and activities resembling those in neurons, and by integrating fluorescence microscopy and functional assays, we showed this cell line to constitute an experimentally attractive system for developing hypotheses that can subsequently be tested in neurons and neuronal models more commonly associated with PD. Our initial focus was on the role of a-syn in the release and recycling of endosomal vesicles, which serve as a proxy for similarly-sized synaptic vesicles. Our results yielded a consistent view that a-syn in helical forms can bind to intracellular pools of these vesicles in the extended-helix conformation and can also engage vesicles docked at the plasma membrane during the process of exocytosis via its broken-helix form. We further found by immunostaining that a-syn associates with mitochondria when expressed in RBL cells^1^, as has been reported previously in other cultured cells and brain tissue.^32–34^

Although PD has a complex etiology involving genetic and environmental factors that vary with individuals, mitochondrial dysfunction is a consistent central feature.^35, 36^ In familial, or autosomal recessive, forms of the disease, disruption of mitochondrial function arises from mutations in genes encoding mitochondrial quality control, including SNCA, LRRK2, VPS35, PARKIN, and PINK-1.^37–41^ Sporadic, or idiopathic, disease and PD-like symptoms can arise from exposure to agents such as rotenone, paraquat, and 1-methyl-4-phenyl-1,2,3,4-tetrahydropyridine (MPTP/MPP^+^) which inhibit complex I of the mitochondrial electron transport chain.^42^ Similarly, carbonyl cyanide m-chlorophenyl hydrazone (CCCP), which inhibits oxidative phosphorylation to produce ATP by uncoupling the mitochondrial membrane potential, has been commonly used to study effects related to PD.^1, 43, 44^ Our previous studies in RBL cells showed that a-syn association with mitochondria increases markedly after treatment with CCCP, and this colocalization is prevented by mutations in either the first (A30P) or second (V70P) N-terminal helices.^1^ Multiple pathological effects have been reported for overexpression of a-syn and variants on mitochondria in model and neuronal systems. These include disruption of mitochondrial fission/fusion, generation of reactive oxygen species (ROS), impaired Ca^2+^ uptake, and reduced ATP production.^34–36, 45, 46^

Mitochondrial stability and function depend on regulated Ca^2+^ flux, and the ER has been reported as the main source of environmentally-stimulated mitochondrial Ca^2+^ uptake in yeast^47^ and mammalian cells,^48^ including neurons.^49^ Functionally, the level of Ca^2+^ in mitochondria regulates the tricarboxylic acid cycle to yield necessary ATP production^50^, and operative sources of Ca^2+^ may differ in the neuronal cell body and in axonal boutons during an action potential.^49, 51^ For most cells and conditions studied, uptake of mitochondrial Ca^2+^ appears to occur mainly at the ER– mitochondria (ER-mito) contact sites, where local Ca^2+^ reaches high concentration levels.^52^ Other critical metabolic functions such as phospholipid and cholesterol exchange also occur in these regions.^53^ Ca^2+^ transfer complexes include several proteins localized to the contact sites, including the voltage-dependent anion-selective channel (VDAC) in the mitochondrial outer membrane; the mitochondrial Ca^2+^ uniporter (MCU) with tissue-specific regulatory proteins (MICU)^54^ reside in the mitochondrial inner membrane. Mitochondria-associated ER membranes (MAM) at the at ER-mito contacts contain the inositol 1,4,5-trisphosphate receptors (IP3R) through which Ca^2+^ is released from the ER, flowing through VDAC to MCU/MICU complex. MAM also contain vesicle-associated membrane protein-associated protein B (VAPB), which serves as a tether by binding to mitochondrial protein tyrosine phosphatase-interacting protein 51 (PTPIP51).^55^

Considering our evidence for stress-related association of a-syn with mitochondria^1^ and previous reports that a-syn can localize to MAM,^32^ we proceeded to investigate more directly the participation of a-syn in modulating Ca^2+^ fluxes into mitochondria, including specific structural determinants and disruptions that may have pathological impact in PD. We employed our established RBL cell model, as well as other cell types used previously in studies related to a-syn and PD: human embryonic kidney (HEK293) cells, murine neuronal 2a (N2a) cells, and differentiated dopaminergic N2a cells. Each of these cell types has little or no endogenous expression of a-syn, and we found that ectopic expression of human a-syn leads to an increase in stimulated mitochondrial Ca^2+^ uptake. We systematically evaluated this effect in RBL cells with a- syn variants, including mutations disrupting a-syn helical structure and the C-terminal region. Our results indicate that membrane-binding by the helix-1 and helix-2 regions of the protein is required, suggesting that a-syn may bridge between mitochondrial and ER membranes. Consistent with this possibility, our high resolution micrographs confirm increases in ER-mito contacts in the presence of a-syn.

We tested effects of mitochondrial stressors CCCP and the MPTP metabolite MPP^+^, which have been related to PD. After recovery from this stress, cells exhibit enhancement of stimulated mitochondrial Ca^2+^ uptake, but this recovery is dramatically impeded in cells expressing a-syn. This inhibition of recovery depends primarily on the unstructured C-terminal tail of a-syn. Toxin-induced mitochondrial stress also causes increased phosphorylation of Ser129, which is accompanied by aggregation of a-syn. Both inhibition of recovery and a-syn aggregation are eliminated when S129 is mutated to alanine, suggesting that these two processes are linked. Overall, our results suggest that a-syn can participate as a tethering protein to modulate Ca^2+^ flux between ER and mitochondria, with potential physiological significance. A-syn can also prevent cellular recovery from toxin-induced mitochondrial dysfunction, possibly in an aggregation-dependent manner, pointing to one pathological mechanism for the role of a-syn in the etiology of PD.

## RESULTS

### A-syn enhances stimulated mitochondrial Ca^2+^ uptake in model RBL mast cells

We showed previously that a-syn colocalizes with mitochondria, especially under conditions of stress.^1^ Because a-syn, mitochondrial dysfunction, and disruption of Ca^2+^ homeostasis are all strongly implicated in PD, we proceeded to evaluate effects of a-syn on a key mitochondrial function: stimulated uptake of Ca^2+^. We designed a fluorescence assay in which RBL cells, transfected with the mitochondrial Ca^2+^ indicator, mito-GCaMP6f^51^ and wildtype a-syn (Wt-syn in pcDNA vector), were sensitized with immunoglobulin E (IgE) and then stimulated by sub-optimally low doses of antigen (0.5-2 ng/ml).

Under these sub-optimal conditions, control cells (transfected in parallel with mito-GCaMP6f and empty pcDNA vector) undergo limited stimulated mitochondrial Ca^2+^ uptake (20% normalized response, averaging over all cells), but cells expressing Wt-syn exhibit a much higher uptake (75%, on average) (Figure 2a, red data and “X”, right axis). This enhanced level is similar to the stimulated response with an optimal dose of antigen for control cells. In evaluating these data for RBL cells, we found that stimulated mitochondrial Ca^2+^ uptake typically occurs in some cells and not others, such that changing the conditions causes the fraction of cells responding to change. By this means of accounting, 33% of the control cells respond and 88% of cells expressing Wt-syn respond (Figure 2a, blue bars, left axis). Because RBL cells consistently exhibited a bimodal response in this assay, we assessed statistical significance with a non-parametric model (Figure 2a, blue P-values). In contrast, HEK293 and N2a cells show normally distributed responses in this same assay (see below). For simplicity, we generally compare responses using values averaged over all cells (e.g., Figure 2a, red “X”, right axis).

**Figure 2.**
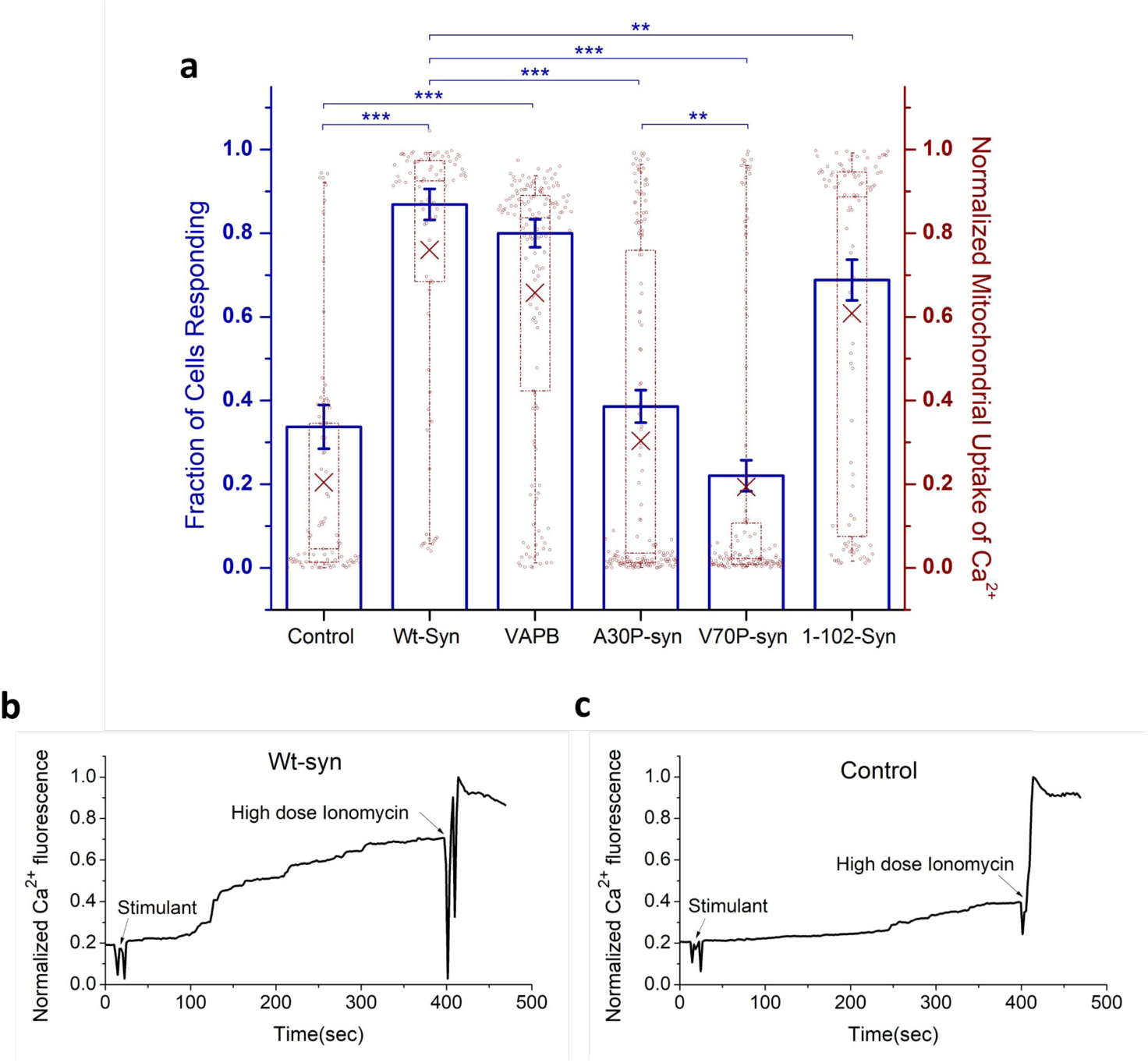
Wt-syn enhances Ag-stimulated mitochondrial Ca^2+^ uptake. RBL cells were co-transfected with plasmids for mito-GCaMP6f and one of pcDNA (empty vector control), VAPB (known ER/mitochondria tether), Wt-syn, or indicated mutant of a-syn. Cells were imaged by confocal microscopy, and mitochondrial Ca^2+^ uptake was stimulated by Ag (DNP-BSA, 1 ng/ml). Mito GCaMP6f fluorescence was monitored in confocal movies before and after stimulation, and after indicated addition of high-dose ionomycin 300-400 sec later to achieve the maximal fluorescence value. ***a***) Left axis: RBL cells exhibit a bimodal distribution of Ag-stimulated responses, and fraction of cells with more than 20% mitochondrial Ca^2+^ uptake (normalized as described in Materials and Methods) is represented by height of blue box; error bars are ± SEM. Statistical significance is based on this fraction; *** represents P-values <0.001, ** represents P-values <0.01. Right axis: Mitochondrial Ca^2+^ uptake for all individual cells evaluated as represented by data points; maroon box plot shows 25th-75th percentile of the data; midline shows median, and X shows average. Data sets are the same for right and left axes: Each sample comprising n∼80 co-transfected cells are from three independent experiments. ***b, c***) Representative traces of mito-GCaMP6f fluorescence integrated over 5-7 cells within confocal fields. Arrows indicate addition of Ag (stimulant) and high dose ionomycin for cells expressing Wt-syn (*b*) or empty pcDNA plasmid (control) (*c*). These traces were extracted from movies similar to those available in Supplemental Materials (Movie S1 for (*b*) and Movie S2 for (*c*)).

Possible mechanisms for the enhancing effect of a-syn relate to the capacity of this protein to bind to membranes in extended or broken helical forms. The latter form has been proposed to tether between different membranes such as synaptic vesicles and the plasma membrane.^21, 56–58^ We hypothesized that a-syn can tether mitochondria to ER, which serves as a source of Ca^2+^. VAPB, an ER protein, is known to act as tether between ER and mitochondria and to enhance stimulated mitochondrial Ca^2+^ uptake by complexing with mitochondrial protein PTPIP51.^59, 60^ We also found that transfection of VAPB into RBL cells causes enhancement in stimulated uptake of Ca^2+^ (average 67%) over control cells (average 20%), very similar to the increase observed with cells expressing Wt-syn (average 75%) (Figure 2a). This result suggests that a-syn can increase Ca^2+^ flow from ER to mitochondria similarly to VAPB, by tethering these two organellar membranes.

To evaluate structural features of a-syn involved in enhancing stimulated Ca^2+^ uptake, we tested variants with mutations in each of the two a-syn helices that are posited to enable tethering of membranes (Figure 1). We found previously that proline point mutations within either helix-1 (A30P) or helix-2 (V70P) locally disrupt the helical structure as determined by NMR measurements, and also disrupt membrane binding (A30P, V70P) or tethering (V70P) capacity of a-syn, as measured with liposome/micelle binding in vitro and stimulated exocytosis of recycling endosomes in cells.^1, 31^ The stimulated mitochondrial uptake assay showed that both mutations significantly attenuate the enhancing effect of Wt-syn: Control empty vector (20%) ≈ V70P-syn (20%) < A30P- syn (30%) << Wt-syn (75%) (Figure 2a). This trend in functional effects correlates with previously observed effects of these mutations on a-syn localization to mitochondria: Wt-syn co-localizes with mitochondria much more strongly than A30P-syn, and V70P-syn co-localization is undetectable.^1^

Because the helix-forming, N-terminal segment (aa 1 to 97 of 140 total) provides the lipid binding affinity of a-syn, we tested whether this segment is sufficient for observed functional effects. We made a C-terminal truncation variant (1-102-syn), which includes aa 1 to 97 and five additional C-terminal residues to ensure an intact helix-2.^61^ Interestingly, 1-102-syn enhances stimulated mitochondrial Ca^2+^ uptake (average 62%), at a level greater than the Control (average 20%) although somewhat less than for Wt-syn (average 75%) (Figure 2a). Together, these results indicate that the N-terminal helices 1 and 2 are primarily involved in mitochondrial binding and tethering capacity of Wt-syn, although the disordered C-terminal residues 103-140 may participate to some extent.

### A-syn enhances stimulated mitochondrial Ca^2+^ uptake in HEK293 cells and N2a cells

HEK293 cells have been widely used as a model system for various cellular pathways, including those implicated in PD. To assess the generality of the effects of a-syn on mitochondrial Ca^2+^ uptake, we used HEK293 cells expressing the same mitochondrial Ca^2+^ indicator, together with either mRFP (control) or a-syn + mRFP (Wt-syn via multicistronic construct P2a-syn-mRFP). Transfected cells were stimulated with either low-dose ionomycin (0.38 μM) or ATP (100 μM). Wt-syn enhances stimulated mitochondrial Ca^2+^ uptake, compared with controls, for both stimulants: from 30% to 50% for ionomycin (Figure 3a and Supplemental Figure S1) and from 22% to 37% for ATP (Figure S1).

**Figure 3.**
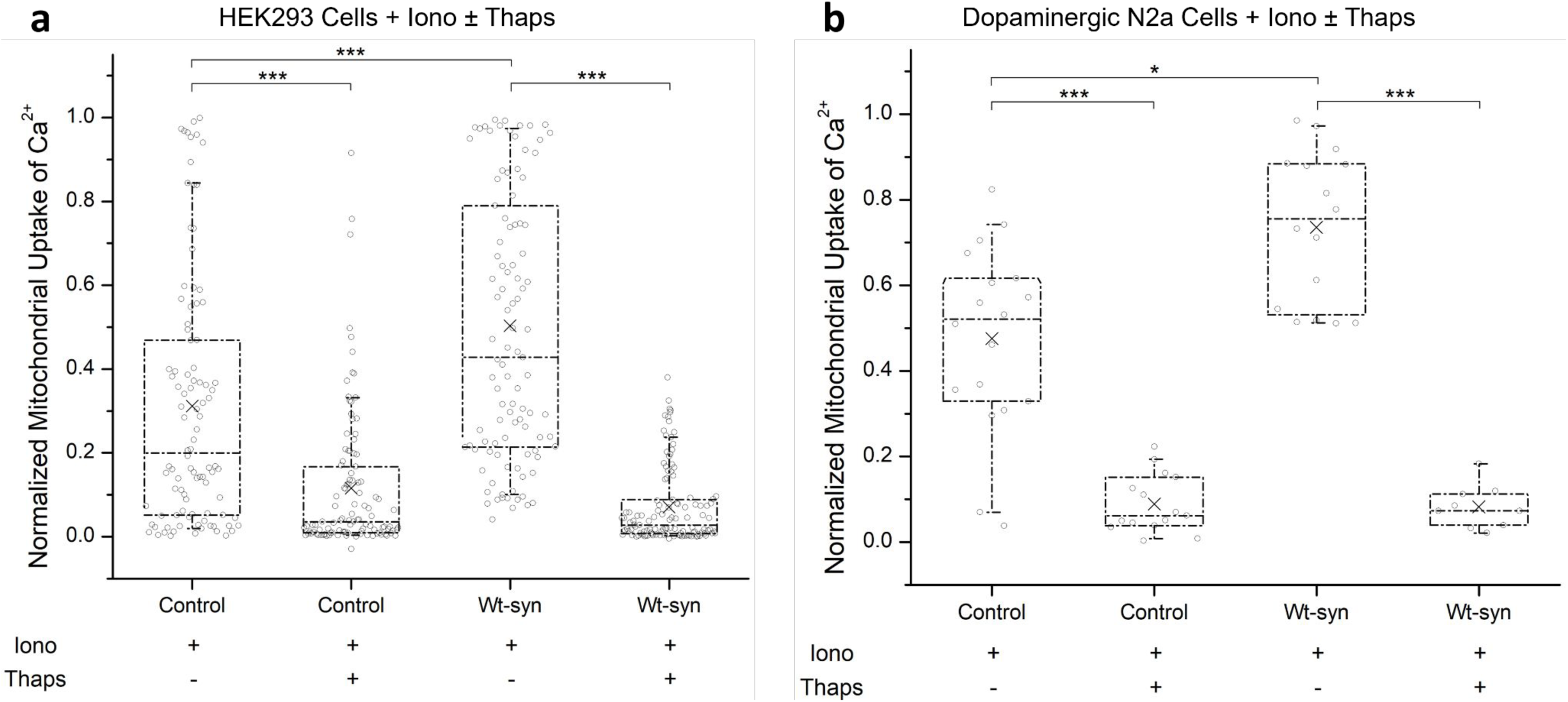
ER is the primary source of Ca^2+^ for stimulated mitochondrial uptake. HEK293 cells (***a***) or dopaminergic N2a cells (***b***) were co-transfected with plasmids for mito-GCaMP6f and either mRFP (control) or Wt-syn (via multicistronic construct syn-p2a-mRFP). Mitochondrial Ca^2+^ uptake was stimulated by low-dose ionomycin; for indicated samples, thapsigargin was added to deplete ER Ca^2+^ stores prior to addition of stimulant. Mito-GCaMP6f fluorescence was monitored in confocal movies before and after stimulation, and after subsequent addition of high-dose ionomycin, several hundred seconds later, to achieve maximal fluorescence. Each sample comprising n∼80 (HEK293) or n∼25 (Dopaminergic N2a) co-transfected cells are from three independent experiments. Data points represent stimulated mitochondrial Ca^2+^ uptake (normalized) for each cell under conditions specified for each sample in Materials and Methods. Box plot shows 25th-75th percentile of the data, midline shows median, and X shows average; *** represents P-values <0.001, * represents P- values <0.05. Representative traces of mito-GCaMP6f fluorescence corresponding to each of the samples in (*a*) and (*b*) are provided in Supplemental Figures S1 and S2, respectively.

Given that PD features selective death of dopaminergic neurons in the substantia nigra region of the brain, we also examined a neuronal cell line, N2a, derived from the mouse neural crest that has been extensively used to study neuronal differentiation, axonal growth, and signaling pathways. We modified previously-established protocols to differentiate N2a cells into dopaminergic neurons, which exhibit morphological neurites and increased levels of tyrosine hydroxylase.^62, 63^ We found that endogenous expression of a-syn in this cell line, before and after differentiation, is below our limits of detection by immunostaining or Western blot. Differentiated N2a cells were transfected with mRFP (control) or Wt-syn (via syn-p2a-mRFP vector), together with the mitochondria Ca^2+^ indicator, followed by stimulation with low-dose ionomycin. Under these conditions, we observed that stimulated Ca^2+^ uptake increases from an average of 48% for control cells to 74% for cells expressing Wt-syn (Figures 3b and S2a,b). Enhancement of stimulated mitochondrial uptake by Wt-syn in HEK293 cells and dopaminergic N2a cells (Figure 3) is consistent with that observed for RBL cells (Figure 2), showing the generality of this effect, which evidently depends on mitochondrial membrane binding/tethering by Wt-syn.

### Endoplasmic reticulum is the main source of Ca^2+^ for stimulated mitochondrial uptake in RBL, HEK293, and N2a cells

Stimulation of cells typically involves release of Ca^2+^ from ER stores, which then triggers opening of plasma membrane channels and Ca^2+^ entry from the extracellular medium into the cytoplasm.^51, 60^ We first considered the possibility that a-syn enhances stimulated mitochondrial Ca^2+^ uptake by increasing ER release of Ca^2+^ into the cytoplasm, and that increase in [Ca^2+^]_cyt_ is required for subsequent uptake into mitochondria. We monitored Ca^2+^ changes in both the mitochondria (mito-GCaMP6f) and cytoplasm (GCaMP3) in RBL cells stimulated with antigen in Ca^2+^-free buffer. Under these experimental conditions, any increase in cytoplasmic Ca^2+^ would come primarily from the ER. We found that expression of Wt-syn enhances stimulated mitochondrial Ca^2+^ uptake (trend similar to Figure 2), but the stimulated change in cytoplasmic Ca^2+^ does not differ significantly in cells expressing Wt-syn compared to control cells (Figure S3). These results are consistent with the view that the stimulated increase in mitochondrial Ca^2+^ comes through a direct route from the ER, rather than indirectly from the bulk pool of cytoplasmic Ca^2+^.

As a more definitive test for the source of Ca^2+^ for mitochondrial uptake, we carried out an experiment similar to Chakrabarti et al^60^ who used the SERCA inhibitor thapsigargin to deplete ER Ca^2+^ stores. They found that pre-treatment of U2OS cells with thapsigargin causes a significant decrease in mitochondrial Ca^2+^ uptake stimulated by ionomycin or histamine. In similar experiments, we tested HEK293 cells (transfected or not with Wt-syn) using thapsigargin to empty ER stores prior to stimulation of mitochondrial Ca^2+^ uptake with ionomycin. The response in HEK293 cells treated with thapsigargin is highly reduced compared to untreated cells: cells expressing Wt-syn decrease from 50% to 10%, and control cells decrease from 30% to 10% (Figures 3a and S1). We carried out the same type of experiment with dopaminergic N2a cells, depleting ER stores with thapsigargin prior to stimulating with ionomycin and observed the same effect on mitochondrial Ca^2+^ uptake: cells expressing Wt-syn decrease from 74% to 8%, and control cells decrease from 48% to 9% (Figures 3b and S2). Thus, for both HEK293 and dopaminergic N2a cells, thapsigargin treatment causes a substantially reduced response in both control cells and cells expressing Wt-syn, indicating that a-syn mainly affects the Ca^2+^ entering mitochondria from ER as opposed to the cytoplasm or other sources. These results are consistent with those from RBL cells, highlighting a-syn-mediated enhancement of stimulated mitochondrial uptake of Ca^2+^, likely by tethering membranes of the two organelles to facilitate direct flow of Ca^2+^ from the ER.

### A-syn enhances ER-mitochondria contacts in cells as quantified in super resolution images

We used structured illumination microscopy (SIM) for ultrastructural evidence of a-syn tethering capacity and consequent impact on ER-mitochondria (ER-mito) contacts. SIM enables both high resolution images^64, 65^ and sample sizes sufficient for effective statistical analysis. N2a cells were transfected with plasmid DNA for ER (STIM1-mApple) and mitochondria (mEmerald-TOMM20) markers and also for Wt-syn (or empty vector for control). Harvested cells were fixed and immunostained with anti-syn antibody to identify cells expressing Wt-syn. The SIM images provide a detailed view of the tubular structure of ER overlapping with mitochondria (Figure 4a,b). N2a cells expressing Wt-syn form more extensive ER-mito contacts compared to control (Figure 4a,b magnified boxes). We observed differences both in the number of contacts and in the length of each individual contact. We calculated values for Pearson’s correlation coefficient (PCC) to determine the overlap of the mitochondrial label with the ER label. This quantification represents the proximity of the two types of organelles and reflects the relative degree of ER-mito contacts in compared samples. The results from analyzing about 35 N2a cells for each condition, shows a significant increase in averaged PCC value from 0.40 to 0.52 (22% increase) for cells transfected with Wt-syn (Figure 4c). This comparison further supports the hypothesis that a-syn acts as a tether between ER and mitochondria, which facilitates an increase in stimulated mitochondrial Ca^2+^ uptake.

**Figure 4.**
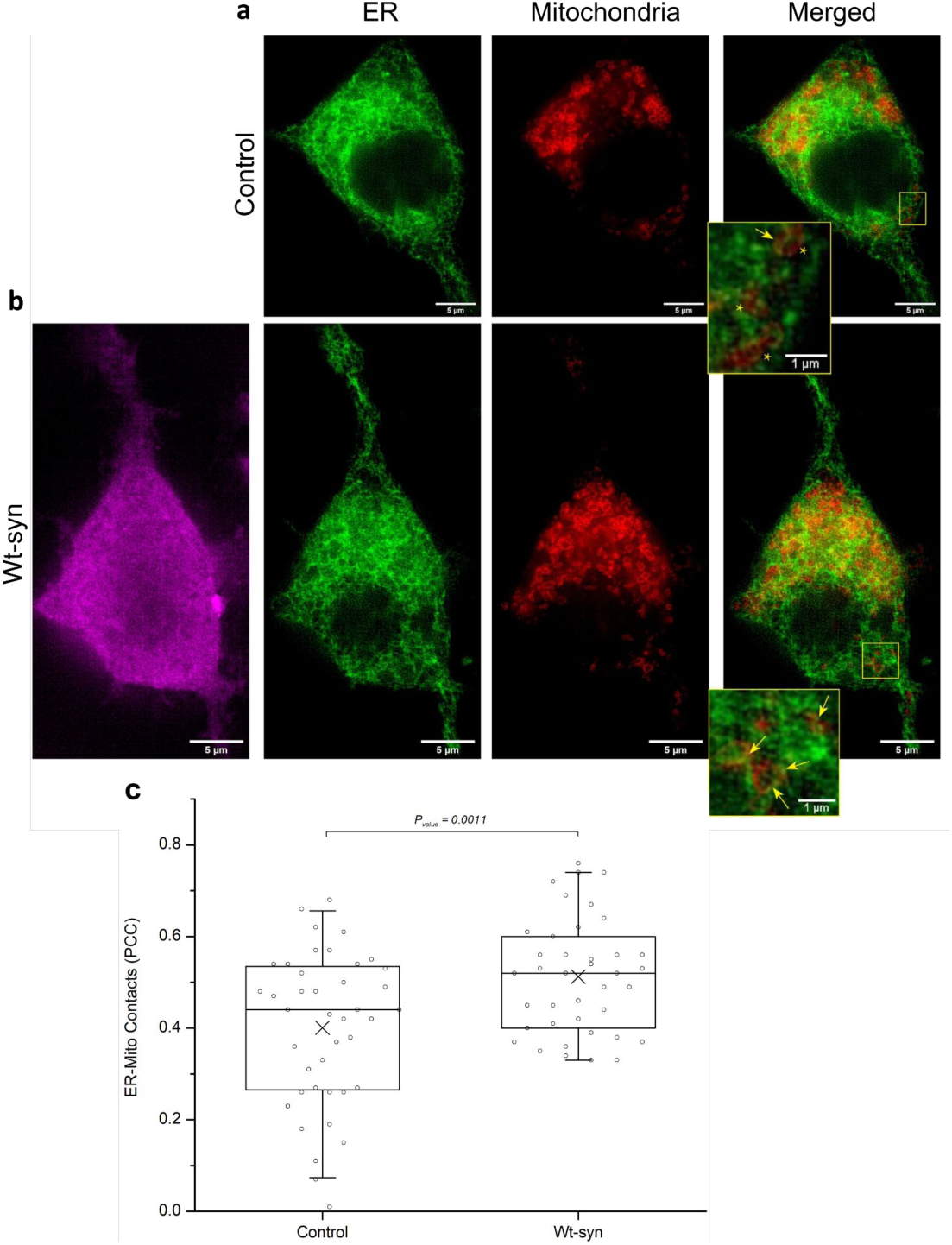
Wt-syn increases contacts between ER and mitochondria. N2a cells transfected with plasmids for pcDNA (control) (***a***) or Wt-syn (***b***), together with fluorescent constructs for both STIM1 (shown as green; ER membrane) and TOMM20 (shown as red; mitochondrial outer membrane) were washed and fixed prior to immunostaining Wt-syn (magenta) for visualization at super resolution with structured illumination microscopy (SIM). Representative images are shown; scale bar = 5μm. Insets in ER/mitochondria-merged images are higher magnification (scale bar = 1μm): Arrows point to regions with clearly contacting ER/mitochondrial membranes; stars label proximal organelle membranes without strong contacts. ***c***) Averaged Pearson’s correlation coefficients (PCC) for mitochondrial membranes overlapping ER membranes were calculated for Wt-syn and control samples as represented in box plot (n∼35 cells). Box shows 25th-75th percentile of the data, midline shows median, and X shows average.

### A-syn disrupts mitochondrial recovery from stress caused by the mitochondrial toxin CCCP in RBL and N2a cells

To investigate possible pathogenic roles of a-syn in mitochondrial dysfunction related to PD, we designed an assay in RBL cells that quantifies the recovery capacity of mitochondria. Brief treatment with CCCP to induce mitochondrial stress was followed by incubation in standard culture medium to facilitate cell recovery. Then, recovery was evaluated by measuring stimulated mitochondrial Ca^2+^ uptake. We showed previously that treatment with CCCP, which has been used in studies of mitochondrial function related to PD^66^, markedly increases colocalization of a-syn with mitochondria, depending on intact N-terminal helices.^1^ In the current study, we found that after acute incubation with CCCP stimulated mitochondrial Ca^2+^ uptake is disrupted (either no response or an erratic response; data not shown). However, after the recovery incubation, control (no a-syn) RBL cells show a level of stimulated mitochondrial Ca^2+^ uptake that is enhanced compared to the response with no CCCP treatment (Figure 5a; Cntrl +CCCP *>* Cntrl -CCCP), indicating a robust compensation mechanism during recovery from stress. This recovery level of enhancement for control cells is similar to that observed for RBL cells expressing Wt-syn with no CCCP treatment (Figure 5a; Cntrl +CCCP ≈ Wt-syn -CCCP). This raises the possibility that the CCCP-recovery mechanism of the control cells involves an increase of mitochondria/ER contacts to enhance stimulated Ca^2+^ transport, and that a similar increase occurs in the absence of CCCP exposure when Wt-syn is expressed. In contrast, RBL cells expressing Wt-syn and treated with CCCP followed by the recovery incubation show a substantially reduced level of stimulated mitochondrial Ca^2+^ uptake (Figure 5a; Wt-syn +CCCP *<* < Cntrl +CCCP ≈ Wt-syn -CCCP). Thus, it appears that Wt-syn interferes with the recovery mechanism operative in control cells.

**Figure 5.**
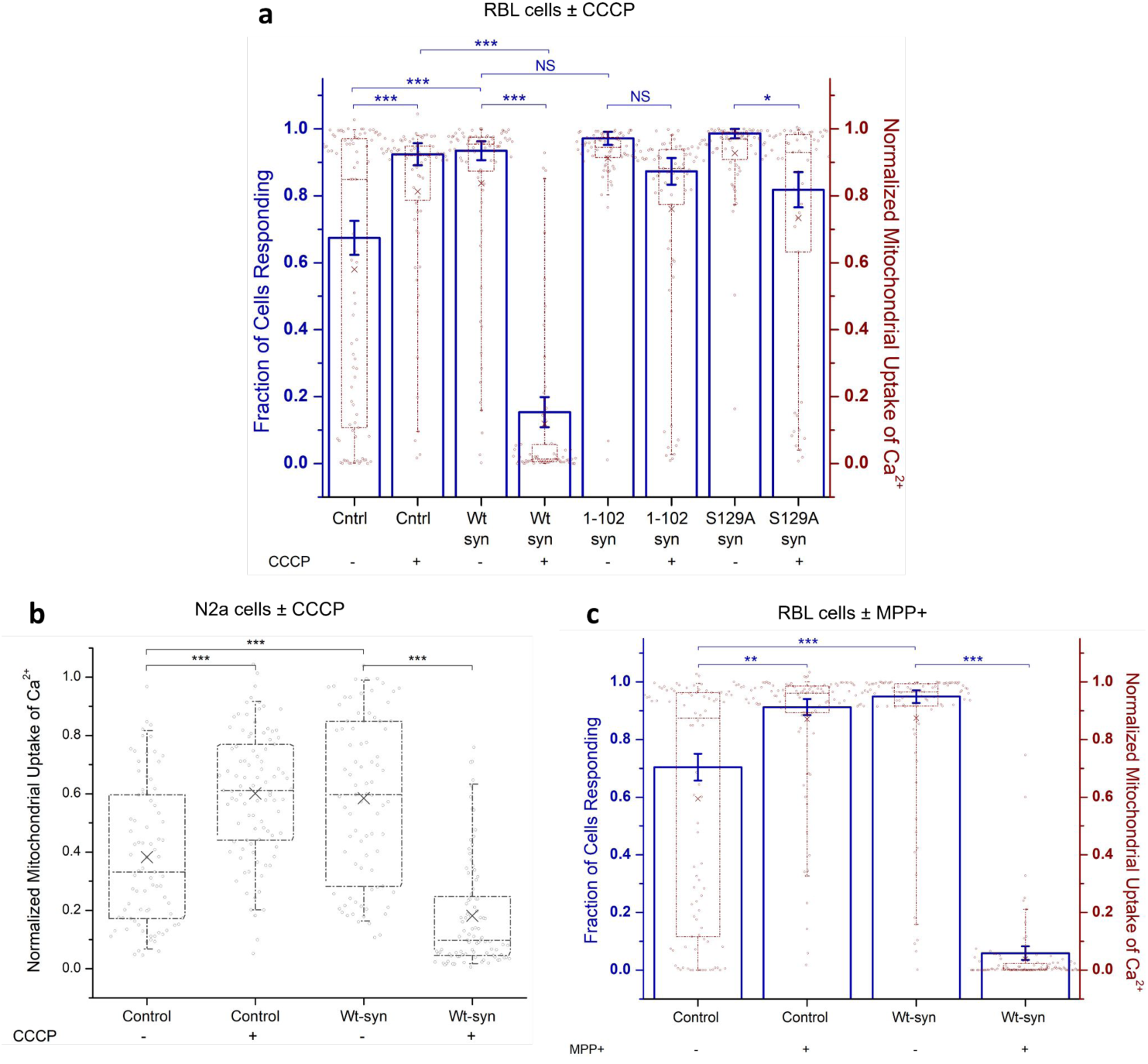
Wt-syn disrupts mitochondrial recovery from stress caused by toxins as measured by stimulated Ca^2+^ uptake. RBL cells (*a, c*) were co-transfected with plasmids for mito-GCaMP6f and one of empty pcDNA plasmid (control), Wt-syn, 1-102-syn (C-terminal truncation), or S129A-syn (eliminated phosphorylation site in C-terminus). N2a cells (*b*) were co-transfected with plasmids for mito-GCaMP6f and either mRFP (control) or Wt-syn + mRFP (Syn-p2a-mRFP). ***a***) RBL cell samples were treated (or not, as indicated) with CCCP, then washed and incubated in fresh media several hours, prior to confocal imaging of mitochondrial Ca^2+^ uptake stimulated by Ag. Left axis: as for Figure 2, cells with more than 20% mitochondrial Ca^2+^ uptake are represented by height of blue bar; statistical significance is based on this fraction; error bars are SEM. Right axis: mitochondrial Ca^2+^ uptake for all individual cells (data points); maroon box shows 25th-75th percentile of the data; midline shows median, and X shows average. Data sets are same for left and right axes (n=70 cells). ***b***) N2a cell samples were treated (or not, as indicated) with CCCP, then washed and incubated in fresh media for several hours, prior to confocal imaging of mitochondrial Ca^2+^ uptake stimulated by low-dose ionomycin. Normalized response for each cell represented by individual points (n=88 cells); box shows 25th-75th percentile of the data, midline shows median, and X shows average. ***c***) RBL cell samples were treated (or not, as indicated) with MPP+, then washed and incubated in fresh media for several hours, prior to confocal imaging of mitochondrial Ca^2+^ uptake stimulated by Ag. Left and right axes are the same as described for *a* (n=100 cells). Details of conditions for all cell samples (*a, b, c*) are specified in Materials & Methods; data sets come from three independent experiments in each case. Statistical significance: *** represents P-values <0.001, ** represents P-values <0.01, * represents P-values <0.05, NS represents not significant.

We also tested the effects of CCCP on basal (non-stimulated) levels of mitochondrial Ca^2+^ after the recovery incubation, and we observed a significant basal increase for control cells (Figure S4; basal Cntl +CCCP > basal Cntrl -CCCP). Interestingly, cells expressing Wt-syn exhibit similarly increased basal mitochondrial Ca^2+^ in the absence of CCCP (Figure S4; basal Wt-syn -CCCP > basal Cntrl -CCCP), again suggesting that control cells may modulate mitochondria/ER contacts in response to CCCP stress and that WT-syn induces the same effect in the absence of CCCP. Accordingly, basal levels do not increase further when WT-syn is present after CCCP treatment and recovery (Figure S4; basal Wt-syn +CCCP ≈ basal Wt-syn -CCCP).

We carried out similar experiments on stimulated mitochondrial Ca^2+^ uptake in N2a cells, testing recovery from CCCP treatment, and we observed the same trends as for RBL cells. Control N2a cells show an enhanced level of stimulated uptake compared to the response with no CCCP treatment (Figure 5b; Control +CCCP *>* Control -CCCP), again indicating a robust compensation mechanism during recovery. The enhancement for control cells is again similar to that observed for N2a cells expressing Wt-syn with no CCCP treatment (Figure 5b; Control +CCCP ≈ Wt-syn - CCCP). In contrast, N2a cells expressing Wt-syn and treated with CCCP followed by the recovery incubation show a substantially reduced level of stimulated mitochondrial Ca^2+^ uptake (Figure 5b; Wt-syn +CCCP *<* < Cntrl +CCCP ≈ Wt-syn -CCCP). A-syn interferes with the recovery mechanism operative in N2a cells, and it is reasonable to infer that the mechanistic and structural underpinnings are similar to those for RBL cells.

### The C-terminal tail of a-syn participates in inhibition of mitochondrial stress recovery

Our finding that Wt-syn inhibits stimulated mitochondrial Ca^2+^ uptake after recovery from chemical stress, in stark contrast to enhancing uptake in the absence of stress, suggests differential structural contributions. To test directly whether the N-terminal, helix-forming region of a-syn is sufficient for abrogation of recovery, we evaluated the C-terminal truncation mutant 1-102-syn (Figure 1) in our assay. We found that RBL cells expressing 1-102-syn recover their stimulation capacity after CCCP stress, similarly to control cells (Figure 5a; 1-102-syn +CCCP ≈ Cntrl +CCCP), with recovery significantly enhanced compared to cells expressing Wt-syn (Figure 5a; 1-102-syn +CCCP >> Wt-syn +CCCP). In contrast, in the absence of CCCP stress, 1-102-syn expression increases the stimulated mitochondrial Ca^2+^ uptake similarly to Wt-syn (Figure 5a; 1-102-syn -CCCP ≈ Wt-syn -CCCP and also Figure 2a). Comparing basal levels of mitochondrial Ca^2+^ we found that cells expressing Wt-syn or 1-102-syn behave similarly to each other and different from control cells (Figure S4; basal Wt-syn ±CCCP ≈ basal 1-102-syn ±CCCP > basal Cntrl -CCCP). These results indicate that the C-terminal tail of a-syn is involved in inhibiting recovery of stimulated Ca^2+^ uptake after mitochondrial stress, whereas the N-terminal helices of a-syn are primarily involved in increasing basal or stimulated Ca^2+^ transport from ER to mitochondria in the absence of stress.

The C-terminal tail is the target of post translational modifications associated with PD.^29, 67–70^ Prominent among these is phosphorylation of Serine-129 (S129) which affects the aggregation state of a-syn as well as interactions with other proteins.^30^ We mutated S129 to alanine and tested this variant (S129A-syn) in our mitochondrial stress recovery assay. Cells expressing S129A-syn exhibit stimulated mitochondrial Ca^2+^ uptake at an enhanced level compared to control cells, similar to cells expressing Wt-syn or 1-102-syn (Figure 5a; S129A-syn -CCCP ≈ Wt-syn -CCCP ≈ 1-102- syn -CCCP > Cntrl -CCCP). After recovery from exposure to CCCP, cells expressing S129A-syn exhibit an enhanced stimulated response, whereas this response is reduced for cells expressing Wt-syn (Figure 5a; S129A-syn +CCCP >> Wt-syn +CCCP). Thus, phosphorylation of S129 may be involved in damaging effects in cells expressing Wt-syn as represented by poor recovery from CCCP-induced mitochondrial stress. A small reduction in the recovery of cells expressing S129A- syn (Figure 5a; S129A-syn +CCCP < S129A-syn -CCCP ≈ Cntrl +CCCP ≈ 1-102-syn +CCCP), leaves open the possibility that residues in the C-terminal tail in addition to S129 contribute to the reduced recovery response.

### A-syn disrupts recovery of mitochondrial stress caused by MPP+ neurotoxin

MPTP, which is physiologically metabolized to the neurotoxin MPP+, is known to cause symptoms of PD in humans and other primates by damaging dopaminergic neurons in the substantia nigra.^71–73^ Used extensively to evaluate mechanisms of PD in model systems^42^, MPP+ has been shown to disrupt mitochondrial function by inhibiting oxidative phosphorylation and consequent ATP production. We found recovery of RBL cells from MPP+ exposure and effects of Wt-syn to be very similar to those we observed for CCCP. Control cells (no a-syn) recovered from MPP+ treatment show an enhanced level of stimulated mitochondrial Ca^2+^ uptake compared to the response with no MPP+ (Figure 5c; Control +MPP+ *>* Control -MPP+), whereas cells expressing Wt-syn have substantially reduced recovery (Figure 5c; Wt-syn +MPP+ *<*< Wt-syn -MPP+). These results are again consistent with the existence in cells of recovery mechanisms to counter effects of PD-related mitochondrial toxins, that are disrupted by a-syn.

### Chemical stress of mitochondria is accompanied by phosphorylation of S129 and aggregation of a-syn

Given the potential involvement of S129 phosphorylation in preventing cellular recovery after mitochondrial stress (Figure 5a) we used confocal fluorescence microscopy and immunostaining to quantify phosphorylated Serine-129 (anti-pSer129-syn; Figure 6) in RBL and N2a cells. Fluorescence intensities observed with each specific antibody were normalized by the expression level of a-syn. CCCP treatment followed by recovery results in increased phosphorylation of S129 in RBL cells transfected with WT-syn (Figure 6b), compared to cells that were not exposed to CCCP (Figure 6a). As expected, RBL or N2a cells transfected with S129A-syn showed negligible fluorescence using anti-pSer129-syn (data not shown) because this variant lacks the phosphorylation site. Quantified over many RBL cells transfected with Wt-syn, the normalized fluorescence of anti-pSer129-syn increases from 0.28 to 0.80 (on average) after recovery from CCCP stress (Figure 6c). In N2a cells, we also observed an increase from 0.22 to 0.48 (Figure 6d). We obtained consistent results in RBL cells treated with MPP+: Anti-pSer129-syn fluorescence increases from 0.37 to 0.95 after recovery from toxin exposure (Figure 6e). These results indicate that phosphorylation of S129 results from exposure to mitochondrial toxins CCCP and MPP+, persisting beyond washing out of the toxin and a recovery incubation.

**Figure 6.**
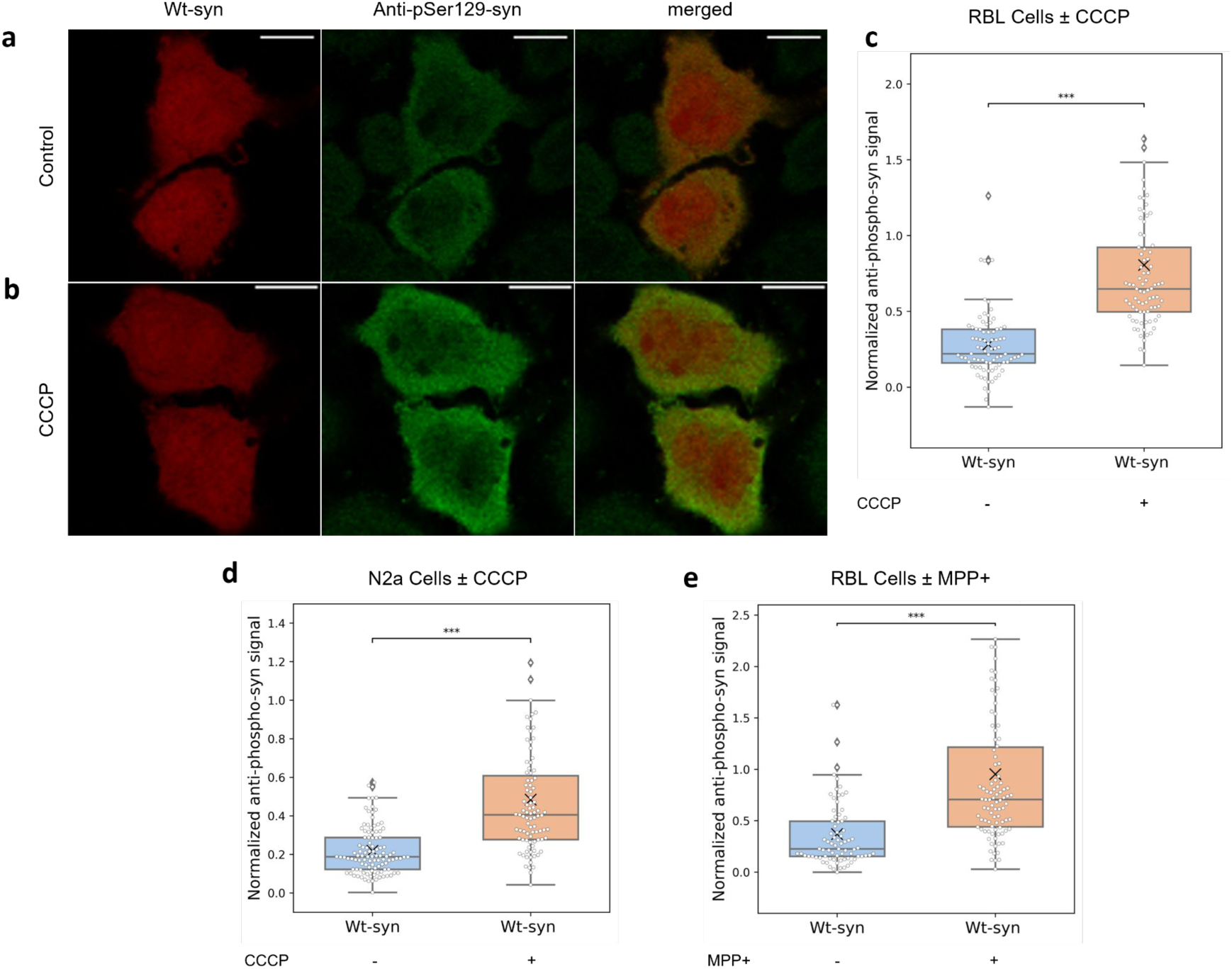
Phosphorylation of Ser129 in Wt-syn increases after toxin treatment and recovery. RBL cells (*a, b, c, e*) and N2a cells (*d*), transfected with plasmid for Wt-syn, were treated, or not, with CCCP (*a, b, c, d*) or MPP+ (*e*), then washed and incubated in fresh media several hours (recovery), prior to immunostaining with anti-pSer129-syn and confocal imaging. ***a, b***) Representative images of RBL cells not treated (control) (*a*) or treated with CCCP (*b*) prior to recovery incubation, showing relative intensities of Wt-syn (red; mRFP from p2a vector) and phosphorylated Wt-syn (green; anti pSer129-syn immunostain). ***c***) RBL cells ± CCCP, quantification of images (n=82 cells for each sample from three independent experiments); pSer129 intensity was normalized by Wt-syn intensity as described in Materials & Methods. ***d***) N2a cells ± CCCP, quantification of images similar to *a* and *b* as described for *c* (n=100 cells for each sample from three independent experiments). ***e***) RBL cells ± MPP+, quantification of images similar to (*a*) and (*b*) as described for (*c*) (n=86 cells for each sample from two independent experiments). Box plots in (*c* – *e*) show 25th-75th percentile of the data; midline shows median, and X shows average. Statistical significance: *** represents P-values <0.001.

Because phosphorylation of S129 is associated with pathological aggregation of Wt-syn^30, 68^, we evaluated the aggregation state of Wt-syn and S129A-syn transfected into RBL and N2a cells. RBL cells immunostained with anti-syn-agg show that CCCP treatment followed by recovery results in increased aggregation of transfected Wt-syn (Figure 7b), compared to cells not exposed to CCCP (Figure 7a). Quantified over many RBL cells transfected with Wt-syn, the normalized fluorescence of anti-syn-agg increases significantly under conditions of CCCP stress (Figure 7c). In contrast, no significant difference in anti-syn-agg staining is detected for RBL cells transfected with S129A-syn with or without exposure to CCCP (Figure 7d-f), indicating that aggregation of Wt-syn in these circumstances requires the presence of Serine-129 and possibly its phosphorylation. We observed the same trends for neuronal N2a cells: Staining by anti-syn-agg increases after exposure to CCCP for cells expressing Wt-syn (Figure 7g) but not for cells expressing S129A-syn (Figure 7h). Use of the neurotoxin MPP+ in RBL cells yielded consistent results: Staining by anti-syn-agg increases after recovery from exposure to MPP+ for cells expressing Wt-syn (Figure 7i) but not for cells expressing S129A-syn (Figure 7j). Together the results shown in Figures 6 and 7 provide structural correlations for the functional outcomes shown in Figure 5, suggesting that a-syn prevents recovery from exposure to mitochondrial toxins by a mechanism involving phosphorylation of S129 and associated aggregation.

**Figure 7.**
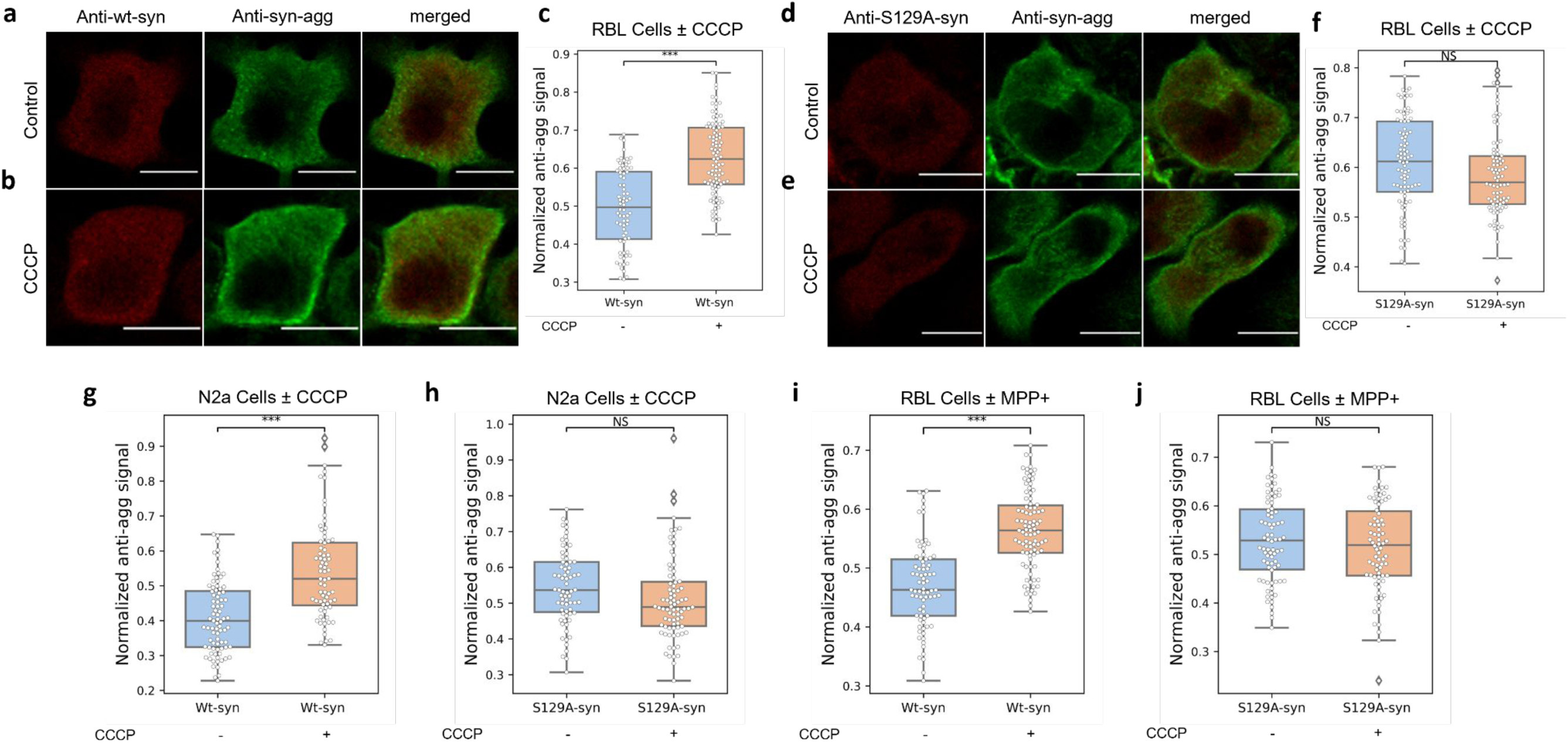
Aggregation of Wt-syn but not S129A-syn increases after toxin treatment and recovery. RBL cells (*a, b, c, d, e, f, i, j*) and N2a cells (*g, h*), transfected with plasmid for Wt-syn or S129A- syn, were treated (or not, as indicated) with CCCP (*a, b, c, d, e, f, g, h*) or MPP+ (*i, j*), then washed and incubated in fresh media for several hours (recovery), prior to immunostaining with antibodies specific for Wt-syn (anti-wt-syn) and aggregated Wt-syn (anti-syn-agg) and confocal imaging. ***a, b***) Representative images of RBL cells expressing Wt-syn and not treated (control) (*a*) or treated with CCCP (*b*) prior to recovery incubation, showing relative intensities of Wt-syn (red) and aggregated Wt-syn (green). **c**) Multiple images of RBL cells expressing Wt-syn ± CCCP similar to (*a*) and (*b*) are quantified (n=85 cells for each sample from three independent experiments); aggregated Wt-syn intensity is normalized by Wt-syn intensity as described in Materials & Methods. ***d, e***) Representative images of RBL cells expressing S129A-syn and not treated (control) (*d*) or treated with CCCP (*e*) prior to recovery incubation, showing relative intensities of S129A-syn (red) and aggregated S129A-syn (green). **f**) Multiple images similar to (*d*) and (*e*) of RBL cells expressing S129A-syn ± CCCP (n=88 cells for each sample from three independent experiments) are quantified; aggregated-Wt-syn intensity was normalized by Wt-syn intensity as for (*c*). ***g***) N2a cells expressing Wt-syn ± CCCP; quantification of multiple images similar to (*a*) and (*b*) (n=78 cells for each sample from three independent experiments) as described for (*c*). ***h***) N2a cells expressing S129A-syn ± CCCP; quantification of multiple images similar to (*d*) and (*e*) (n=77 cells for each sample from three independent experiments) as described for *f*. ***i***) RBL cells expressing Wt-syn ± MPP+; quantification of images similar to (*a*) and (*b*) (n=86 cells for each sample from two independent experiments) as described for (*c*). ***j***) RBL cells expressing S129A-syn ± MPP+; quantification of images similar to (*d*) and (*e*) (n=79 cells for each sample from two independent experiments) as described for (*f*). Box plots in (*c*), (*f*), and (*g – j*) show 25th-75th percentile of the data; midline shows median. Statistical significance: *** represents P-values <0.001; *NS* represents not significant.

## DISCUSSION

Two cardinal features of PD are disruptions in mitochondrial function and dynamics^35, 46, 74^ and aggregation and deposition of the protein a-syn. These features may intersect in the etiology of PD, as a-syn has been associated with disruption of normal mitochondrial functions, including dysregulated fission/fusion, generation of reactive oxygen species (ROS), impaired Ca^2+^ uptake, and reduced ATP production. ^34–36, 45, 46, 75–78^ Mitochondrial Ca^2+^ levels are intimately linked to mitochondrial function, and their regulation is known to be important for ATP production^79^, control of cell death^80^, ROS signaling^81^, and buffering of cytosolic Ca^2+^ levels.^82^ A-syn has also been implicated in the regulation of cellular Ca^2+^ homeostasis,^83–86^ including within mitochondria. As part of an earlier study, we reported that a-syn association with mitochondria is enhanced under conditions of mitochondrial stress.^1^ These initial results motivated our current study to investigate the effects of a-syn in environmentally-stimulated mitochondrial Ca^2+^ uptake. We first examined RBL cells, which we previously established as a versatile system for evaluating a-syn interactions with membranes associated with specific cell functions, such as release and recycling of endosomal vesicles approximating synaptic vesicles.^1^ We further corroborated our observations in cell lines HEK293, and N2a, which are commonly used as models for neurons in PD and other studies. Our findings extend and provide new mechanistic insights to previous evidence that a-syn can modulate the flow of Ca^2+^ from ER to mitochondria.

### A-syn facilitates stimulated mitochondrial Ca^2+^ uptake from ER depending on N-terminal helices and bridging across organelles

With IgE-sensitized RBL cells we showed that ectopically expressed, human a-syn (Wt-syn) increases mitochondrial Ca^2+^ uptake stimulated by antigen, and the level of this stimulated increase is similar to that caused by over-expression of ER protein VAPB (Figure 2). Miller and colleagues have shown in neurons and neuronal cell lines that VAPB localizes in the MAM sub-compartment of ER membrane, which is enriched with negatively charged phospholipids and cholesterol.^87^ VAPB binding to mitochondrial protein PTPIP51 serves to tether these two organelles in contact sites to facilitate flow of Ca^2+^ from ER stores through IP_3_ receptors to VDAC-MCU channels in the mitochondria.^60, 88^ The increased concentration of Ca^2+^ sequestered in the contact sites is sufficient for uptake by the low affinity MCU/MICU complex.^54^ Our results suggest that, similar to VAPB, a-syn can serve as a tether to add or further tighten ER-mito contact sites, thereby enhancing Ca^2+^ sequestration and increasing stimulated mitochondrial Ca^2+^ uptake from ER stores. Our analysis of SIM images showing increased ER-mito contacts in the presence of Wt-syn (Figure 4) further supports this interpretation. Guardia-Laguarta et al. reported the presence of a-syn in MAM fractionated from cultured cell models overexpressing a-syn and from normal human/mice brain tissues.^32^ Supported by structural studies on model and cell membranes^1, 21, 22, 56– 58, 89^ this tight association is consistent with dual-anchor a-syn attachment to ER and mitochondria membranes, possibly under both physiological and pathological conditions.

Our results showing a-syn enhancement of basal and stimulated mitochondrial Ca^2+^ uptake by increasing ER-mito contacts can be compared to previous reports.^83, 90–92^ In a study on SH-SY5Y human neuroblastoma cells and HeLa cells, Cali et al showed that expression or knock-down of a- syn lead to an increase or decrease, respectively, in stimulated mitochondrial Ca^2+^ uptake and in ER-mito contacts^90^ in agreement with our results. Two other studies in SH-SY5Y and HEK293 cells reported that overexpression of a-syn reduces stimulated mitochondrial Ca^2+^ uptake by interfering with ER-mito tethering by either VAPB-PTPIP51^92^ or IP3R-GRP75 ^93^ interactions and thereby reducing ER-mito contacts.^92^ In these studies, rounding of mitochondria^92^ or mitochondrial fragmentation and sensitization to depolarization^93^ accompanied a-syn over-expression, suggesting that expression levels were high enough to cause mitochondrial stress. This could explain the discrepancy with our own work, and may indicate that the authors were instead observing effects of a-syn aggregation related to our observations after treatment with mitochondrial toxins. In support of this view, a subsequent study by Cali et al.^94^ showed that higher levels of a-syn results in a loss of enhanced Ca^2+^ uptake. More generally, it seems clear that differences in results and/or interpretation among these studies are likely due in part to different levels of a-syn expression and perhaps also to other differences in experimental conditions, underlining challenges in elucidating physiologic and pathologic roles of a-syn. Indeed, both overexpression and loss of a-syn have been linked to mitochondrial dysfunction in mice,^95–98^ although aspects of this too remain controversial. Using a simpler model system, we showed previously that high and low expression levels of a-syn in RBL cells lead to different functional outcomes.^1^

Our focus on structure-function relationships, which we evaluate by introduction of a-syn variants into cells with little or no endogenous a-syn, points to mechanisms by which a-syn may participate in mitochondrial Ca^2+^ homeostasis and disruption. Although intrinsically disordered in solution, a-syn has been shown to adopt an amphipathic helical structure in the presence of negatively charged lipid surfaces^48, 49^, such as located in MAMs. The distribution of broken helix (residues 3-38 and 46-93; Figure 1) *vs* extended helix (residues 3-97) depends on membrane curvature and proximity of a second membrane.^21, 56, 58^ Our previous NMR measurements showed that proline point mutations A30P within helix-1 and V70P within helix-2 locally disrupt the helical structure of the protein^1^, while also reducing the overall affinity of the protein for membranes.^31^ These perturbations have functional consequences. For example, we showed previously that whereas Wt-syn inhibits stimulated exocytosis of recycling endosomes, V70P-syn abrogates this effect, evidently by preventing a-syn bridging between vesicles and the plasma membrane.^1^ In current studies we found that, compared to Wt-syn, A30P-syn expression causes much less enhancement of stimulated mitochondrial Ca^2+^ uptake and that V70P-syn expression shows almost no enhancement (Figure 2). Together with our previous findings,^1^ our results are consistent with a- syn tethering ER and mitochondrial membranes via the broken-helix form to enhance the stimulated flow of Ca^2+^ into the mitochondria.

We also evaluated deletion of a-syn’s disordered C-terminal tail in light of previous reports indicating its role in protein-protein interactions, including interactions with the mitochondrial outer membrane anion channel VDAC^99, 100^, which could contribute to ER-mito tethering. We observed that the 1-102-syn variant only slightly reduces the enhancing effect of WT-syn (Figure 2), indicating the primacy of N-terminal helices in the enhancement of Ca^2+^ uptake. In contrast with our results, a recent study reported that the A30P mutation of a-syn does not abrogate its ability to enhance uptake.^94^ Because the effects we observe for the A30P mutation on bridging-related functions^1^ (Figure 2) are milder than those of the V70P mutation, the A30P variant may retain activity at higher expression levels, suggesting again that different expression levels in the two studies are responsible for the different observations. In addition, Cali et al^90^ found that removal of the C-terminal tail of a-syn by truncation at residue 97 eliminated enhancement of Ca^2+^ uptake, in contrast with our finding that truncation at a-syn residue 102 only mildly reduced this effect. This discrepancy may result from destabilization of a-syn helical structure and membrane binding by truncating the protein at position 97, very close to the C-terminal end of a-syn helix-2^16, 20, 101^ (Figure 1) with consequent loss of enhanced Ca^2+^ uptake.

Our results strongly implicate the ER as the main source of Ca^2+^ transported into mitochondria under our conditions (Figure 3), supporting ER-mito contacts as the relevant context for the effects of a-syn. Our results are consistent with those of Chakrabarti et al^60^ who similarly treated U2OS cells with thapsigargin to empty ER Ca^2+^ stores and demonstrated no significant increase in mitochondrial Ca^2+^ stimulated by ionomycin or histamine, in contrast to non-treated cells. Ashrafi et al^51^ used a similar approach to demonstrate ER as the source of stimulated mitochondrial Ca^2+^ uptake in HEK293 cells. They also showed that axonal mitochondria in hippocampal neurons require brain-specific MCU regulator MICU3 to allow efficient Ca^2+^ uptake from the cytoplasm (and not ER) as necessary for rapid ATP production during action potentials. It may be that mitochondrial uptake depends on Ca^2+^ release from the ER in different subcellular regions in neurons, and ER-dependence has been shown for dendrites of cortical pyramidal neurons.^49^ Close examination with focused ion beam-scanning electron microscopy revealed mito/ER contacts in cell bodies, axons, and dendrites from mouse brain tissue.^64, 102^

### A-syn-mediated susceptibility to damage from mitochondrial toxins is accompanied by phosphorylation of S129 and aggregation

Our finding that a-syn interferes with the capacity of cells to recover from mitochondrial toxins points to a pathway that may contribute to the pathological effects of a-syn. PD arises from and manifests in a complex combination of mitochondrial dysfunctions, including under-production of ATP, over-production of ROS, and mis-regulated Ca^2+^ flow. Using mitochondrial Ca^2+^ uptake as a robust assay, we could test effects of toxins related to PD and participation of a-syn. Along with rotenone and paraquat, MPTP/MPP+ is a known neurotoxin, with limited exposure known to cause Parkinsonian symptoms in humans and other mammals.^46^ The MPTP metabolite MPP+ has been shown to accumulate selectively in dopaminergic neurons and cause their apoptosis.^103^ This class of neurotoxins are Complex 1 inhibitors, and pathology is thought to arise in part from increased levels of ROS. Carbonyl cyanide phenylhydrazones, including CCCP, act to dissipate the mitochondrial membrane potential, and this class also increases ROS through pathways that may include reaction with glutathione.^104^ As shown by others in cells, consequences of increased ROS include phosphorylation of a-syn S129, which can accompany a-syn aggregation.^36^ We previously showed that colocalization of Wt-syn with mitochondria increases markedly after stressing with CCCP.^1^

In the present study we found that acute treatment with CCCP or MPP+ disrupts stimulated mitochondrial Ca^2+^ uptake in RBL and N2a cells. Remarkably, the cells recover this mitochondrial capacity, and stimulated Ca^2+^ uptake is actually enhanced after toxin removal and further incubation in normal media (Figure 5). The similarity of this enhanced uptake to that mediated by VAPB or a- syn in the absence of toxins suggests that the recovery mechanism may involve increased ER-mito contacts. The fact that mitochondria can recover from toxin exposure attests to cellular resilience and robust repair pathways. For example, increases in ROS induced by CCCP results in activation of various protein kinases and phosphatases, in some cases by directly oxidizing the thiol groups of cysteine residues. These and other perturbations initiate signaling pathways of nuclear factor erythroid 2-related factor 2 (Nrf2) and transcription factor EB (TFEB) as part of an integrated cellular stress response.^104^ The increase in basal level mitochondrial Ca^2+^ we observed in control cells after CCCP recovery, compared to no treatment, (Figure S4) is consistent with such induced repair mechanisms.

We discovered that recovery of stimulated mitochondrial Ca^2+^ uptake after toxin exposure is inhibited in RBL and N2a cells expressing Wt-syn (Figure 5), indicating that perturbations mediated by a-syn when these toxins are present compromise stress response mechanisms. In contrast to Wt-syn, 1-102-syn expression allows mitochondrial recovery from toxin exposure, implicating the C terminal tail in impeding recovery. A prominent feature of the a-syn C-terminal tail is phosphorylation of S129, which is intimately linked to PD pathology.^29, 30^ The kinase responsible for S129 phosphorylation is Polo-like kinase 2 (PLK2),^105–108^ which has also been reported to function in mitochondrial Ca^2+^ uptake at ER-mito contacts^109^ and to be up-regulated by mitochondrial stress.^110, 111^ Accordingly, increased S129 phosphorylation under conditions of mitochondrial stress and increased ROS has been established^109^ and has been related to a-syn aggregation, changes in subcellular distributions and neuronal loss in transgenic mice.^36^ We therefore examined phosphorylation of S129 (Figure 6) and a-syn aggregation (Figure 7) in RBL and N2a cells after recovery from exposure to toxins MPP+ and CCCP. Consistent with expected effects of increased ROS due to these toxins, we observed increases in Wt-syn phosphorylation and aggregation, which correlate with abrogated recovery of stimulated mitochondrial Ca^2+^ uptake.

Strikingly, we find that S129A-syn does not inhibit recovery from mitochondrial toxins and does not result in associated aggregation of a-syn, suggesting that S129 phosphorylation is upstream of both effects. Remaining unresolved are the mechanisms and structural underpinnings for inhibition of mitochondrial Ca^2+^ uptake by pS129-syn or its accompanying aggregation. We previously observed that CCCP treatment causes recruitment of a-syn to mitochondria^1^, and this is reasonably the first step involved. Recruitment may result from increased ER-mito contacts that we posit occur as part of the toxin recovery process and which would present a-syn with favorable binding opportunity via broken-helix bridging between the closely apposed ER and mitochondrial membranes. In this model, S129 phosphorylation occurs after a-syn relocates to stressed mitochondria, likely by PLK2, which has increased activity under this stress.^110, 111^ Indeed, accumulation of pS129-syn at damaged mitochondrial membranes has been reported in a PD-related synucleinopathy.^112^

Subsequent steps involved in a-syn aggregation and inhibition of mitochondrial recovery are more difficult to discern, particularly as they relate to each other. S129 phosphorylation has been shown to modulate a-syn interactions with other cellular proteins,^28, 113, 114^ and loss or gain of interactions with particular proteins may underlie pathological consequences. For example, the negatively charged C-terminal tail of a-syn has been reported to inhibit VDAC via electrostatic interactions with its positively charged pore.^99, 100^ The interaction could be enhanced by S129 phosphorylation, providing a potential mechanism for direct interference with recovery of Ca^2+^ uptake at ER-mito contacts. It is possible that S129 phosphorylation promotes a-syn aggregation directly or indirectly through other interactions. While this has remained controversial, with reports differing on whether pS129 promotes or inhibits aggregation,^115^ the reality is likely to be context dependent. Two recent studies reported that a-syn recruitment to mitochondria results in a-syn aggregation^116, 117^ but these studies did not examine the phosphorylation state of S129. Finally, the mechanisms by which aggregates of a-syn may lead to mitochondrial dysfunction, including inhibiting their recovery after toxin exposure, are likely complex^76, 118–121^ and remain a subject of ongoing investigation.

## CONCLUSIONS

Overall, our investigation supports the possibility that a-syn modulates basal and stimulated mitochondrial Ca^2+^ uptake under physiological conditions. We observed that expression of Wt-syn increases mitochondrial Ca^2+^ levels in stimulated RBL cells and neuronal cell models, including N2a cells that are differentiated to mature, dopaminergic neurons similar to those found in the substantia nigra. By comparing structural variants of a-syn and by controlling the sources of Ca^2+^ we provided evidence for Wt-syn acting as a tether to strengthen ER-mito contacts, depending on the integrity of its N-terminal helices. The ultimate functional outcome of this structural interaction may be concentration dependent. Considering the evidence from the literature and our results, we suggest a concentration dependent role for wt a-syn in modulating mitochondrial Ca^2+^ uptake. Within a physiological concentration range the enhancing effect of wt a-syn on mitochondrial Ca^2+^ uptake may help maintain mitochondrial Ca^2+^ homeostasis before and after stimulation. However, for concentrations above the physiological range, the tethering capacity of wt a-syn to increase contacts between ER and mitochondria and/or its aggregation may cause pathological effects on mitochondrial function and morphology.

After treatment with mitochondrial toxins, including MPTP/MPP^+^ known to induce parkinsonism, we directly observed pathological effects of a-syn. We showed that Wt-syn disrupts the recovery of mitochondria from toxin-induced stress and consequently causes severe inhibition of stimulated mitochondrial Ca^2+^ uptake. The unstructured C-terminal tail of a-syn participates in causing this damage, which is accompanied by and may depend on phosphorylation of S129, and also results in a-syn aggregation. The roles of specific and nonspecific structural interactions of a- syn with membranes and other proteins in normal physiology and in disease remain an intriguing puzzle awaiting future investigation of this complex system in cellular and neuronal models and in primary neurons susceptible to PD and other synucleinopathies.

## MATERIALS AND METHODS

### Reagents

1-methyl-4-phenylpyridinium iodide (MPP+ iodide), retinoic acid, dibutyryl cyclic AMP sodium salt, thapsigargin, carbonyl cyanide m-chlorophenyl hydrazine (CCCP), and EGTA were from Sigma-Aldrich (St. Louis, MO). Trypsin-EDTA, 0.2 µm TetraSpeck™ beads, Alexa Fluor 488-, Alexa Fluor 568-, and Alexa Fluor 647-labeled goat anti-mouse or anti-rabbit IgG secondary antibodies were from Invitrogen (Carlsbad, CA; CAT#: A21121, A21124, A21240, A21241, A11034; 1:200 dilution). Transfection reagents TransIT-X2® and Lipofectamine® 2000 were from Mirus Bio (Madison, WI) and Thermo Fisher Scientific (Waltham, MA), respectively. VECTASHIELD HardSet Antifade Mounting Medium was from Vector Laboratories (Burlingame, CA). Mouse monoclonal IgG1 anti-α-synuclein antibodies 3H2897(CAT#: sc-69977; 1:200 dilution), 42/α-Synuclein (CAT#: 610787; 1:200 dilution) and anti α,β-synuclein (AB_2618046; 1:150 dilution) were from Santa Cruz Biotechnology (Dallas, TX), DSHB (University of Iowa) and BD Biosciences (Franklin Lakes, NJ), respectively. These anti-a-syn antibodies were optimized for use across experiments, with the same one used in any given experiment. We found that 42/α-Synuclein was most sensitive for labeling S129A-syn, similarly to Wt-syn. Monoclonal anti-tyrosine hydroxylase antibody (Product ID: 22941; 1:220 dilution) was from ImmunoStar (Hudson, WI). Recombinant rabbit monoclonal anti phosphorylated-α-synuclein (Ser129) antibody (Clone#: EP1536Y; 1:400 dilution) and anti-α- synuclein aggregate antibody (Clone#: MJFR-14-6-4-2; 1:1000 dilution) were acquired from Abcam (Cambridge UK).

### Cell Culture

RBL-2H3 cells were cultured as monolayers in minimal essential medium (Invitrogen) with 20% fetal bovine serum (Atlanta Biologicals, Atlanta, GA) and 10 µg/ml gentamicin sulfate (Invitrogen) as previously described.^122^ HEK293 (a gift from Barbara Hempstead, Weill Cornell Medicine) and Neuro2a (N2a from ATCC) cells were cultured in Dulbecco’s modified eagle medium (DMEM, Invitrogen) with 10% fetal bovine serum and 50 µg/ml of Pen-Strep (Invitrogen). Adherent cells were harvested by treatment with Trypsin-EDTA (0.05%) for 8-10 min (RBL-2H3 cells) or 2-3 min (HEK293, N2a cells), 3-5 days after passage.

### Cell expression plasmids

The cDNA for cell expression of human Wt-syn in pcDNA 3.0 vector was a gift from Dr. Chris Rochet (Purdue University). Other plasmids for expression of human a-syn mutants (A30P-syn, V70P-syn, 1-102-syn and S129A-syn) were created within this pcDNA vector by site directed mutagenesis using Phusion High-Fidelity DNA Polymerase (New England Biolabs)^1^. The plasmid for GCaMP3 was purchased from Addgene (# 22692), and those for mito-GCaMP6f, mito-jRCaMP1b, and ER-GCaMP6f were gifts from Dr. Tim Ryan (Weill Cornell Medicine).^51^ The plasmid for VAPB was a gift from Dr. Chris Steffan (University College, London). The plasmid for STIM1-mApple was prepared in our lab as described previously^123^ and that for mEmerald-TOMM20-N-10 was a gift from Michael Davidson (http://n2t.net/addgene:54282 ; RRID:Addgene_54282).

Syn-p2a-mRFP is a multicistronic vector encoding Wt-syn and mRFP simultaneously, allowing Wt-syn expression level to be determined without adding a tag to the relatively small a-syn protein. To create the syn-p2a-mRFP plasmid, the cDNA encoding human Wt-syn was introduced into a vector from Clontech (Mountain View, CA) containing the mRFP sequence, using Hind III and KpnI restriction sites, followed by insertion of the p2a sequence using KpnI and XmaI restriction sites.^124^ The p2a motif was prepared by annealing the following primers:

p2a-syn-mRFP forward: CGGGAAGCGGAGCTACTAACTTCAGCCTGCTGAAGCAGGCTGGAGACGTGGAGGAGAACCCTGGACCTGC

p2a-syn-mRFP reverse: CCGGGCAGGTCCAGGGTTCTCCTCCACGTCTCCAGCCTGCTTCAGCAGGCTGAAGTTAGTAGCTCCGCTTCCCGGTAC

### Experiments with RBL cells

#### Transfection by electroporation

RBL-2H3 cells were harvested three to five days after passage, and 5 x 10^6^ cells were suspended in 0.5 ml of cold electroporation buffer (137 mM NaCl, 2.7 mM KCl, 1 mM MgCl_2_, 1 mg/ml glucose, 20 mM HEPES, pH 7.4). Co-transfections used a reporter plasmid DNA (5 µg mito-GCaMP6f), together with 5 µg of Wt-syn in pcDNA 3.0 vector or empty pcDNA 3.0 vector (control), VAPB, or a-syn mutants: A30P-syn, 1-102- syn, V70P-syn). We have found for RBL cells that cells transfected with two constructs express both proportionally, such that a fluorescent construct can be used as an expression reporter for cells co-transfected with a non-fluorescent construct.^1^ Cells were electroporated at 280 V and 950 μF using Gene Pulser X (Bio-Rad), then immediately resuspended in 6 ml medium and cultured in MatTek dishes (2 ml/dish) (MatTek Corporation, Ashland, MA) for 24 hr to recover; the medium was changed after live cells became adherent (1-3 hr). For stimulation by antigen, cells were sensitized with 0.5 μg/ml anti-2,4-dinitrophenyl (DNP) IgE during the recovery period.^125^

We took several measures to ensure consistency of gene expression, in multiple experiments over different days. We visualized the levels of expressed a-syn variants using immunofluorescence imaging labeling with an antibody specific for all tested variants (typically AB_2618046; 1:150 dilution). We also confirmed with regular testing that RBL cells co-transfected with mito-GCaMP6f and Wt-syn show a strong correlation with respect to fluorescence intensity from mito-GCaMP6f compared to immunostained Wt-syn. Therefore, we could evaluate mito GCaMP6f fluorescence as a reliable reference for consistency of transfection efficiency and thereby a measure of the consistency in transfection of a-syn variants. Our previous measurements, including quantitative western blots,^1^ indicate that levels of Wt-syn in RBL-2H3 cells transfected with this amount of Wt-syn plasmid express are around ∼10 μΜ, which is within the physiological concentration range of this protein found in neurons. The observed immunofluorescence is also consistent with this expression level based on our prior experience.^1^

#### Stimulated mitochondrial Ca^2+^ uptake assay

After the electroporation recovery period and prior to imaging, cells were washed once and then incubated for 5 min at 37°C with buffered saline solution (BSS: 135 mM NaCl, 5 mM KCl, 1 mM MgCl_2_, 1.8 mM CaCl_2_, 5.6 mM glucose, 20 mM HEPES, pH 7.4). Then, mito-GCamp6f fluorescence was monitored for 20 sec prior to stimulation with 1 ng/ml DNP-BSA (antigen). After 6-8 min of stimulation, a high dose of ionomycin (5 μM) was added to saturate the mitochondria with Ca^2+^ for normalization of the stimulated fluorescence. Cells were monitored with time by confocal microscopy (Zeiss 710) using a heated, 40X water objective. Mito-GCaMP6f was excited using the 488-nm line of a krypton/argon laser and viewed with a 502- 551 nm band-pass filter. Mitochondrial Ca^2+^ uptake was quantified in individual cells as described below (Equation 2).

#### Mitochondrial stress recovery assay

RBL cells were co-transfected with 5 µg mito GCaMP6f, together with 10 µg of syn-p2a-mRFP, S129A-syn-p2a-mRFP, 1-102-syn, or mRFP (control). Prior to imaging, cells were divided into three groups: Group 1 cells were treated with 375 μM of MPP+ or 10 μM of CCCP in BSS buffer for 30 min, followed by washing with culture medium and incubation for 6 hours (after MPP+ treatment) or 3 hours (after CCCP treatment) in culture medium at 37°C to recover. Group 2 cells were treated with CCCP or MPP+ as for Group 1 cells but did not go through recovery and were washed three times with BSS buffer at 37°C. Group 3 cells (control) were treated as for Group 1 cells, except without addition of any drugs. Mitochondrial uptake of Ca^2+^ was stimulated with 100 ng/ml DNP-BSA after the recovery period for Group 1 and 3 cells and immediately after the drug treatment for Group 2 cells.

#### Immunostaining of a-syn variants

Cells were electroporated with selected plasmids and cultured in MatTek dishes for 24 hr, then fixed with 4% paraformaldehyde + 0.1% glutaraldehyde. After washing, fixed cells were labeled in PBS with 10 mg/ml BSA (PBS/BSA) using a monoclonal anti-a-syn antibody, followed by a secondary antibody conjugated to Alexa Fluor (488 or 568 or 647, depending on experiment), and then imaged by confocal microscopy.

#### Measuring phosphorylation of serine 129 in Wt-syn

Cells were transfected with 10 μg of syn-p2a-mRFP and cultured in MatTek dishes before treating with CCCP or MPP+ or buffer (control) then incubating in culture medium, as described for the mitochondrial stress recovery assay. Cells were fixed, labeled in PBS/BSA using monoclonal anti-pSer129-syn (1:400) followed by secondary antibody conjugated to Alexa Fluor488, and then imaged by confocal microscopy. The microscopy was performed using a Zeiss LSM 880 confocal microscope equipped with 1.4 NA, 40X oil immersion objective and GaAsP detectors, using 488 nm and 561 nm laser lines.

#### Measuring aggregation level of Wt-syn and S129A-syn

Cells were transfected with 15 μg of pcDNA-S129A-syn or pcDNA-Wt-syn and cultured in MatTek dishes before treating with CCCP or MPP+, or not, then incubated in culture medium as described for the mitochondrial stress recovery assay. Cells were fixed, labeled in PBS/BSA using monoclonal anti-a-syn antibody (1:200) or monoclonal anti-a-syn aggregate antibody (1:1000) followed by respective secondary antibody conjugated to Alexa Fluor (488 or 647, respectively). Confocal imaging with a Zeiss LSM 880 microscope was then performed as described above, using 488 nm and 633 nm laser lines.

### Experiments with HEK293 cells

#### Chemical transfection

HEK293 cells were plated at a concentration of 1×10^5^ cells/ml in DMEM and 10% FBS for 48 hours and then transfected with mRFP or Syn-p2a-mRFP (1 μg) and mtio-GCaMP6f (1 μg) using Lipofectamine® 2000 Transfection Reagent according to the manufacturer’s instructions. 24 hours after the chemical transfection and prior to imaging, cells were washed once and then incubated for 5 min at 37°C in BSS. The stimulated mitochondrial Ca^2+^ uptake assay was carried out as described for RBL cells, except that for HEK293 cells, mito-GcaMP6f fluorescence was monitored for 20 sec prior to stimulation with either 100 μM ATP or 0.38 μM ionomycin. After 3-4 min of stimulation, a high dose of ionomycin (5 μM) was added to saturate the mitochondria with Ca^2+^ for fluorescence normalization. In specified experiments, cells were treated with 0.67 mM thapsigargin for 2-3 min before stimulation with 0.38 μM ionomycin. Cells were monitored by confocal microscopy as in the RBL cell assay; in addition, the 561 nm laser line was used to quantify the expression level of syn-p2a-mRFP. The consistency of the results we obtain in HEK293 cells with those we observe in the RBL cells indicates that synuclein expression levels are in the same physiological range as in the RBL cells and that we are not in the overexpression regime, where the effects of a-syn on mitochondrial Ca^2+^ entry are different and opposite to those we observed.^92–94^

### Experiments with N2a cells

#### Cell differentiation

We tested a combination of conditions to differentiate N2a cells into dopaminergic neurons, which are characterized by increased levels of tyrosine hydroxylase and dopamine. N2a cells normally produce low levels of these two components, and these are enhanced in the presence of dibutyryl cyclic adenosine monophosphate (dbcAMP).^62^ Teatment with retinoic acid also drive these cells towards differentiation and formation of neuronal morphological features including neurites. We optimized differentiation by culturing N2a cells (1x 10^5^ cells/mL) in DMEM + 1% FBS with 1 mM of dbcAMP for 48 hours before washing and culturing in DMEM + 1% FBS with10 μM of retinoic acid for 48 hours. We confirmed differentiation by measuring three to four-fold cellular increase in tyrosine hydroxylase levels with immunostaining^62^ and observing morphological features such as increased size, irregular shape and development of neurites. Differentiated N2a cells were washed with fresh medium of DMEM + 10% FBS before the transfection step.

Differentiated N2a cells were transfected with mRFP or syn-p2a-mRFP (1 μg) and mito GcaMP6f (1 μg) using TransIT-X2® Dynamic Delivery System (Mirus Bio) according to manufacturer’s instructions before reculturing in normal medium for 24 hours. Prior to confocal imaging, cells were washed once and then incubated for 5 min at 37°C with BSS. Mito-GcaMP6f fluorescence was monitored for 20 sec prior to stimulation with 2.5 μM ionomycin. After 1-2 min of stimulation, a high dose of ionomycin (8 μM) was added to saturate the mitochondria with Ca^2+^ for fluorescence normalization. In specified experiments, cells were treated with 0.67mM thapsigargin for 2-3 min before stimulation with 2.5 μM ionomycin. Cell fluorescence was monitored by confocal microscopy as in the RBL and HEK cell assays; in addition, the 561 nm laser line was used to quantify the expression level of syn-p2a-mRF. The consistency of the results we obtain in N2a cells with those we observe in the RBL cells indicates that synuclein expression levels are in the same physiological range as in the RBL cells and that we are not in the overexpression regime, where the effects of a-syn on mitochondrial Ca^2+^ entry are different and opposite to those we observed.^92–94^

#### Structured illumination microscopy (SIM)

Undifferentiated N2a cells were plated on collagen coated MatTek dishes at a concentration of 1×10^5^ cells/ml in DMEM and 10% FBS for 48 hours before transfection with selected constructs. ER membranes were labeled with STIM1 conjugated to rapidly maturing monomeric red fluorescent protein mApple (STIM1-mApple). Mitochondria were labeled with Translocase of Outer Mitochondrial Membrane 20 (TOMM20) conjugated with rapidly maturing monomeric green fluorescent protein mEmerald (mEmerald-TOMM20). Cells were transfected with pcDNA or Wt-syn in a pcDNA vector (1 μg), together with STIM1-mApple (1 μg) and mEmerald-TOMM20 (1 μg) using TransIT-X2® Dynamic Delivery System according to manufacturer’s instructions, followed by reculturing for 24 hours. Then, cells were washed with PBS (pH = 7.4) three times, fixed and immunostained for Wt-syn as described for RBL cells to visualize and quantify expression. To improve the signal to noise ratio and minimize photobleaching, the samples were covered with VECTASHIELD HardSet Antifade Mounting Medium and incubated in a hypoxia chamber for 1 hour under 1% oxygen until the medium hardened. Samples were then imaged on a Zeiss Elyra microscope utilizing a 63× oil objective, and 0.2 µM TetraSpeck™ beads were used as fiducial markers to ensure alignment of different imaging channels. Collected images were aligned and processed using the Zeiss Zen software to produce the high-resolution SIM images. The fluorescence levels for all samples in each channel were adjusted within similar thresholds to ensure consistency for the following steps. The red (ER) channel was used as a mask to delineate ER location throughout an individual cell. Then, applying this mask the correlation between the red channel and the green (mitochondria) channel, was quantified using the Pearson correlation coefficient (PCC):

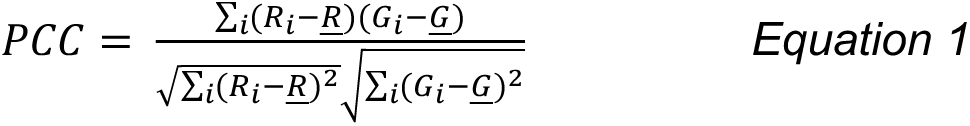

where R represents the signal from the red channel, G represents the signal from the green channel and <u>#x1D445;</u> or <u>#x1D43A;</u> is the mean for the specified signal. About 35 cells were collected for each of control cells and cells expressing Wt-syn over 6 -7 separate days of experiment.

#### Mitochondrial stress recovery assay

Undifferentiated N2a cells were co-transfected with 1 µg mito-GCaMP6f, together with 1 µg of syn-p2a-mRFP (Wt-syn) or mRFP (control). Prior to imaging, cells were divided into three groups: Group 1 cells were treated with 40 μM of CCCP in BSS buffer for 45 min, followed by washing with culture medium and incubation for 7 hours in culture medium at 37°C to recover. Group 2 cells were treated with CCCP as for Group 1 cells but did not go through recovery and were washed three times with BSS buffer at 37°C. Group 3 cells were treated like Group 1 cells but without CCCP and were washed with BSS buffer at 37°C twice. Mitochondrial uptake of Ca^2+^ was stimulated with 2.5 μM ionomycin after the recovery period for Group 1 and 3 cells and immediately after the drug treatment for Group 2 cells.

#### Measuring phosphorylation of serine 129 in Wt-syn

Undifferentiated N2a cells were transfected with 1 μg of syn-p2a-mRFP and cultured in MatTek dishes for 24 hours before treating with CCCP or buffer then incubating in culture medium, as described for the mitochondrial stress recovery assay. Cells were fixed, labeled in PBS/BSA using monoclonal anti-pSer129-syn (1:400) followed by secondary antibody conjugated to Alexa Fluor488, and then imaged by confocal microscopy as described for RBL cells.

#### Measuring aggregation level of Wt-syn

Undifferentiated N2a cells were transfected with 1 μg syn-p2a-mRFP and cultured in MatTek dishes for 24 hours before treating with CCCP or buffer then incubated in culture medium, as described for the mitochondrial stress recovery assay. Cells were fixed, labeled in PBS/BSA using monoclonal anti-a-syn antibody (1:200) and monoclonal anti a-syn aggregate antibody (1:1000) followed by respective secondary antibody conjugated to Alexa Fluor (488 or 647, respectively), and then imaged by confocal microscopy as described for RBL cells.

### Offline image analysis using ImageJ (National Institutes of Health)

#### Stimulated mitochondrial Ca^2+^ uptake assay

Time traces of mito-GCaMP6f fluorescence in individual cells monitored by confocal microscopy were normalized to a 0-1 scale using the following equation:

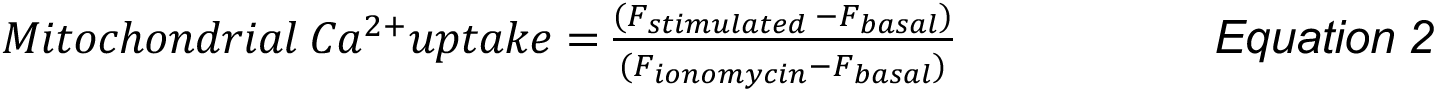

where #x1D439;_-./.0_ is the averaged measured GCaMP6f fluorescence before adding stimulant, #x1D439;_/12340.156_ is the averaged highest fluorescence values after adding stimulant and before high-dose ionomycin addition, and #x1D439;_278739:28_ is the averaged highest steady values following high-dose ionomycin addition. All three averages were calculated over five points to reduce the effects of the random noise.

The resting level for mitochondrial Ca^2+^ in cells was calculated as follows:

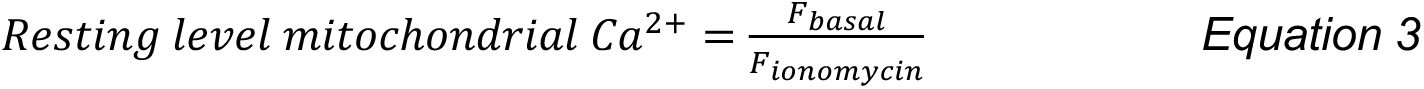

#### Phosphorylation of Ser129 in Wt-syn

Individual cells in fixed confocal images were segmented, and the following fluorescence signals were quantified: mRFP (measure of Wt-syn expression level); anti-pSer129 (Alexa Fluor 488; measure of phosphorylated Wt-syn); weak background signal from five cells not expressing Wt-syn (Alexa Fluor 488 channel). The phosphorylation level was calculated by subtracting the background noise from the anti-pSer129- syn signal and dividing this value by Wt-syn expression level to account for the variability of Wt-syn expression in different cells.

#### Aggregation of Wt-syn and S129A-syn

Individual cells in fixed confocal images were segmented, and the following fluorescence signals were quantified: anti-a-syn antibody (Alexa Fluor 647; measure of Wt-syn expression); anti-a-syn aggregate antibody (Alexa Fluor 488, measure of aggregation level of Wt-syn); weak fluorescence signal averaged over five cells not expressing Wt-syn (Alexa Fluor 488 and 647 channels; background noise). After subtracting the background noise from both fluorescent antibody values, the aggregation level was calculated by dividing the anti-a- syn aggregate signal by Wt-syn expression level to account for the variability of Wt-syn expression in different cells.

### Statistical analyses for cell samples

These were performed with Origin Pro (OriginLab Corp) and Microsoft Excel. For results with normal distribution of data points, statistical significance was determined by a One-Way ANOVA (Analysis of Variance) followed by Tukey’s post hoc test using Origin software. We found that data for RBL cells from assays of stimulated mitochondrial Ca^2+^ uptake (with and without toxin treatment) is not normally distributed and was best interpreted in terms of responding and non-responding cells. In that case, data were binarized based on a reasonable cut off for the non-responding cells. Then the Kruskal-Wallis rank sum test for multiple independent samples was performed followed by Dunn p-values, further adjusted by the Benjamini-

Hochberg FDR method. For both types of statistical analysis, the level of significance is denoted as follows: *P < 0.05, **P< 0.01, ***P < 0.001.

## DATA AVAILABILITY

The fluorescence imaging datasets generated and analyzed as part of this study are available from the corresponding authors on reasonable request.

## ACKNOWLEDGEMENTS

We are grateful to Tapojyoti Das (Weill Cornell Medical College) for helpful discussions. We thank Dr. Tim Ryan (Weill Cornell Medical College) for Ca^2+^ indicator constructs and helpful discussions. Fluorescence imaging was carried out in the Cornell University Biotechnology Resource Center with funding for the Zeiss LSM 710 confocal microscope (NIH S10RR025502) and Zeiss Elyra microscope (NSF 1428922). Research support came from NIH grant R01GM117552 (B.B. and D.H.) and R35GM136686 (D.E.).

## AUTHOR CONTRIBUTIONS

The experiments were designed, executed, and analyzed primarily by M.R., with contributions from A.W.-W. and T.W.. B.B., D.E. and D.H. provided supervision and participated in the design and interpretation of experiments. The manuscript was written by all authors.

## ADDITIONAL INFORMATION

Supplementary information accompanies the paper.

## SUPPLEMENTAL FIGURE LEGENDS

**Figure S1.**
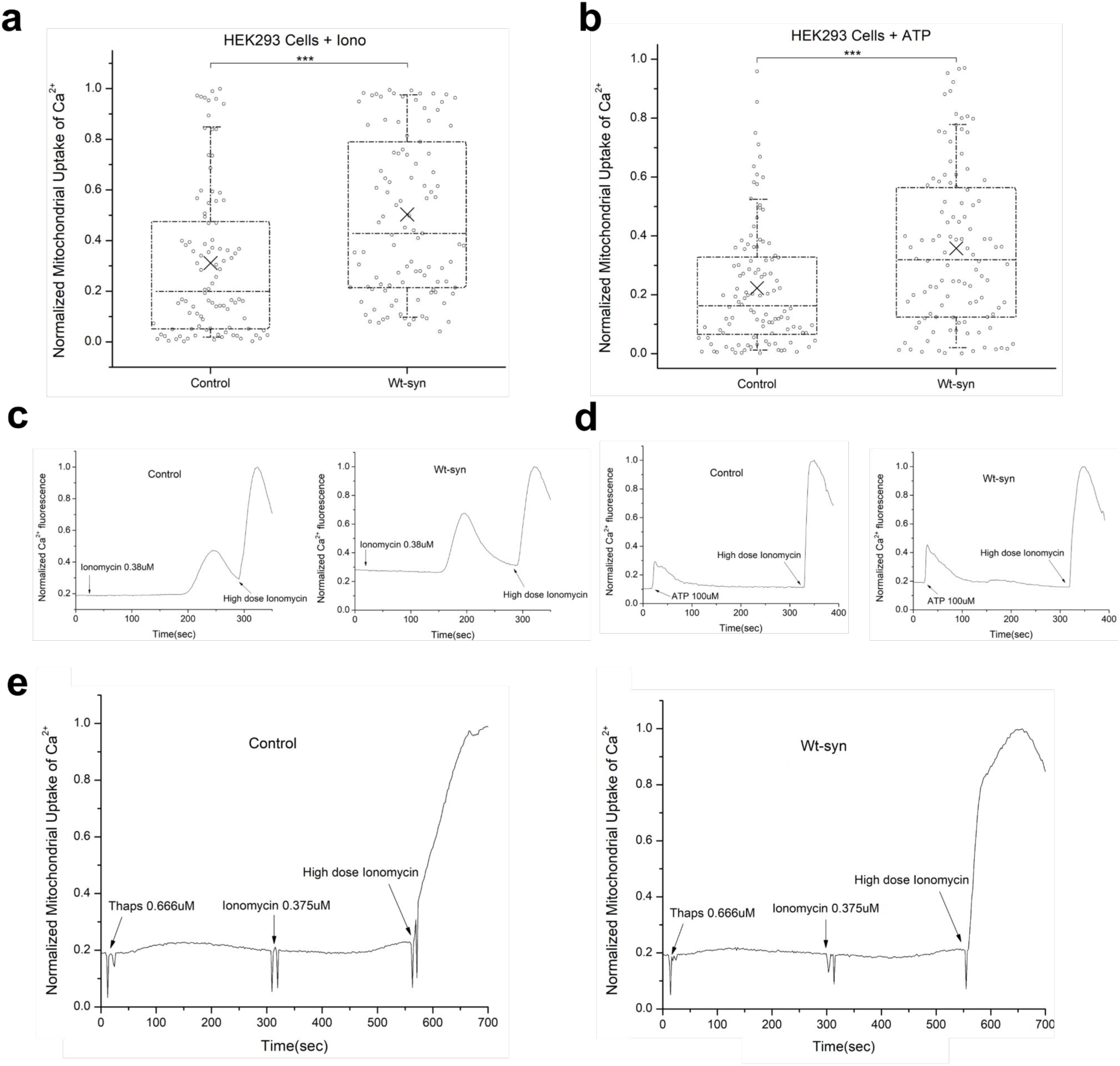
Wt-syn enhances mitochondrial Ca^2+^ uptake from ER in HEK293 cell as stimulated by ionomycin or ATP. Comparative stimulation data and representative traces for experiments shown in Figure 3a of main text. HEK293 cells were co-transfected with plasmids for mito-GCaMP6f and mRFP (control) or Wt-syn (via multicistronic construct Syn-p2a-mRFP). Mitochondrial Ca^2+^ uptake was stimulated by low-dose ionomycin (0.38 μM, *a, c*) or ATP (100 μM, *b, d*). Mito-GCaMP6f fluorescence was monitored in confocal movies before and after stimulation, and after indicated addition of high-dose ionomycin (5 μM, 300-400 sec later) to achieve the maximal fluorescence value. ***a, b***) Averaged mitochondrial Ca^2+^ uptake stimulated by ionomycin (*a*) or ATP (*b*). Each sample comprising n∼80 co-transfected cells are from three independent experiments. The box shows 25th-75th percentile of the data, midline shows median, and X shows average; *** represents P-values <0.001. ***c, d, e***) Representative traces of mito-GCaMP6f fluorescence integrated over 5-7 cells within confocal fields. Arrows indicate addition of thapsigargin (0.67 mM) (*e*), low-dose ionomycin (*c, e*) or ATP (*d*) and high-dose ionomycin (*c, d, e*) for control cells and cells expressing Wt-syn.

**Figure S2.**
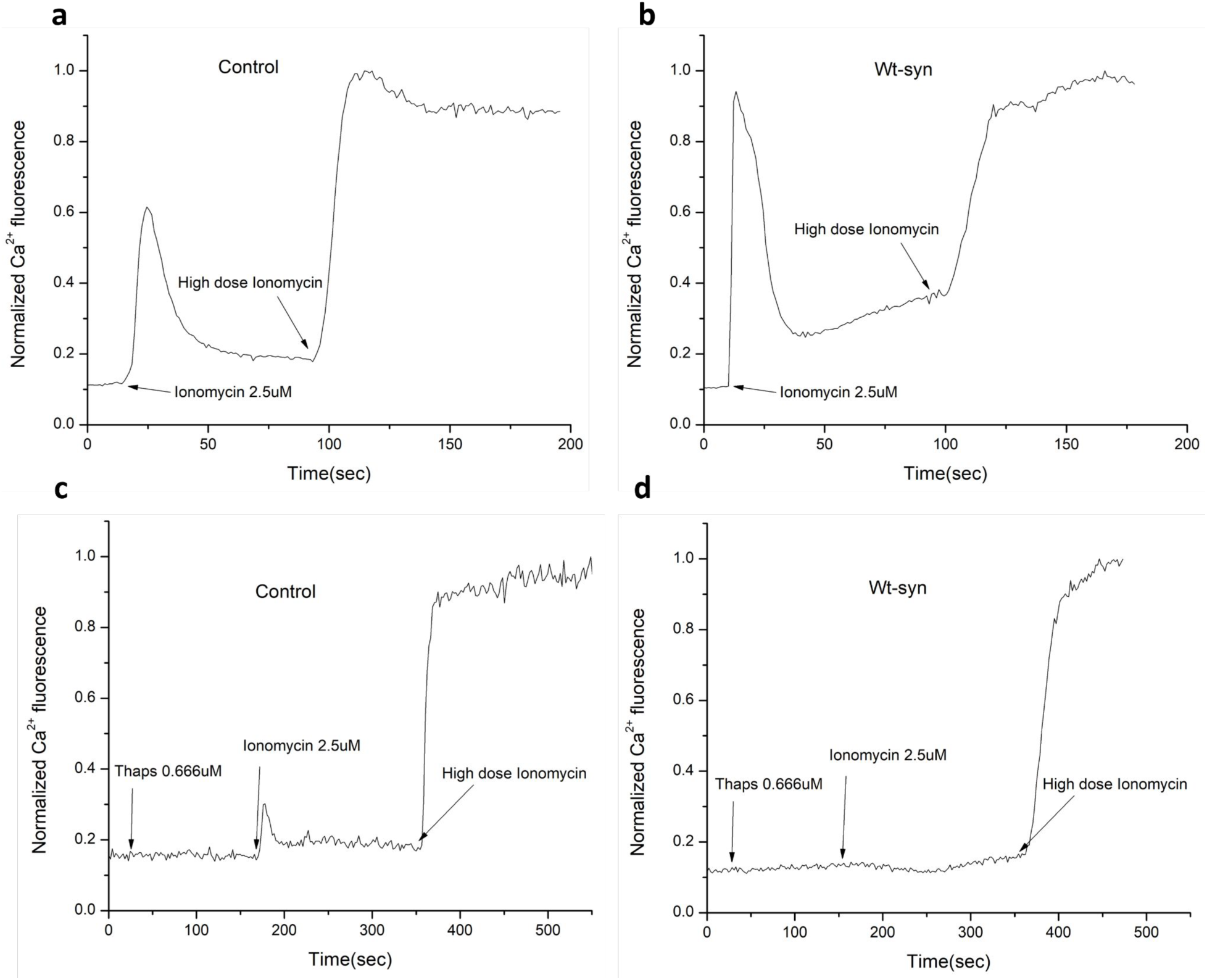
Wt a-syn enhances mitochondrial Ca^2+^ uptake from ER in dopaminergic N2a cells. **a, b, c, d**) Representative traces for experiments shown in Figure 3b of main text. Dopaminergic N2a cells were co-transfected with mito-GCaMP6f and mRFP (control) (*a, c*) or Wt-syn (via multicistronic construct Syn-p2a-mRFP) (*b, d*). Cells were stimulated by low-dose ionomycin (2.5 μM) without (*a, b*) or with (c, d) pre-treatment with thapsigargin (0.67 mM). Mito-GCaMP6f fluorescence was monitored in confocal movies before and after thapsigargin and stimulation, and after addition of high-dose Ionomycin (8 μM, 200-300 sec later) to achieve maximal fluorescence value. Traces integrated from confocal fields containing 1-3 cells.

**Figure S3.**
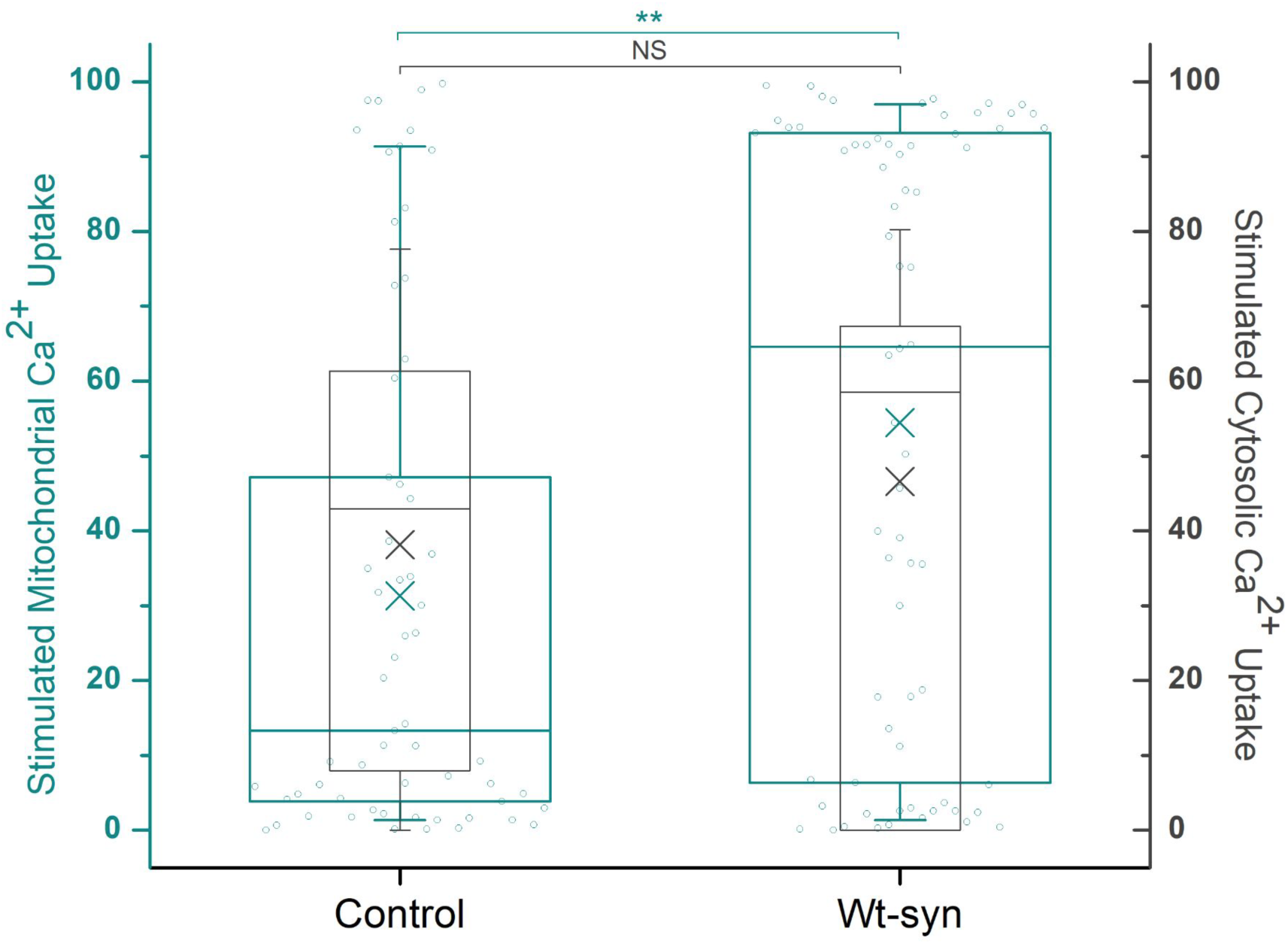
Enhancement of stimulated mitochondrial Ca^2+^ uptake by Wt-syn is not due to increased cytosolic Ca^2+^ uptake. RBL cells were co-transfected with plasmids for either mito-jRCaMP1b (mitochondrial Ca^2+^ indicator), GCaMP3 (Cytosolic Ca^2+^ indicator) and one of pcDNA (empty vector control) or Wt-syn. Harvested cells were transferred to Ca^2+^-free BSS buffer containing 2 μM EGTA, then imaged by confocal microscopy. Mitochondrial Ca^2+^ (left axis) and cytosolic Ca^2+^ (right axis) were monitored in confocal movies before and after stimulation by antigen, and after indicated addition of high-dose ionomycin (5 μM, 300-400 sec later) to achieve maximal fluorescence for each Ca^2+^ indicator. Normalized data points shown are stimulated mitochondrial Ca^2+^ uptake. Normalized data for both mitochondrial (green, left) and cytosolic (black, right) Ca^2+^ increases are represented in superimposed box plots. Each sample comprising n∼60 cells are from three independent experiments. The box shows 25th-75th percentile of the data, midline shows median, and X shows average. Error bars are ± SEM; ** represents P-values <0.01, NS represents not significant (P-values > 0.05).

**Figure S4.**
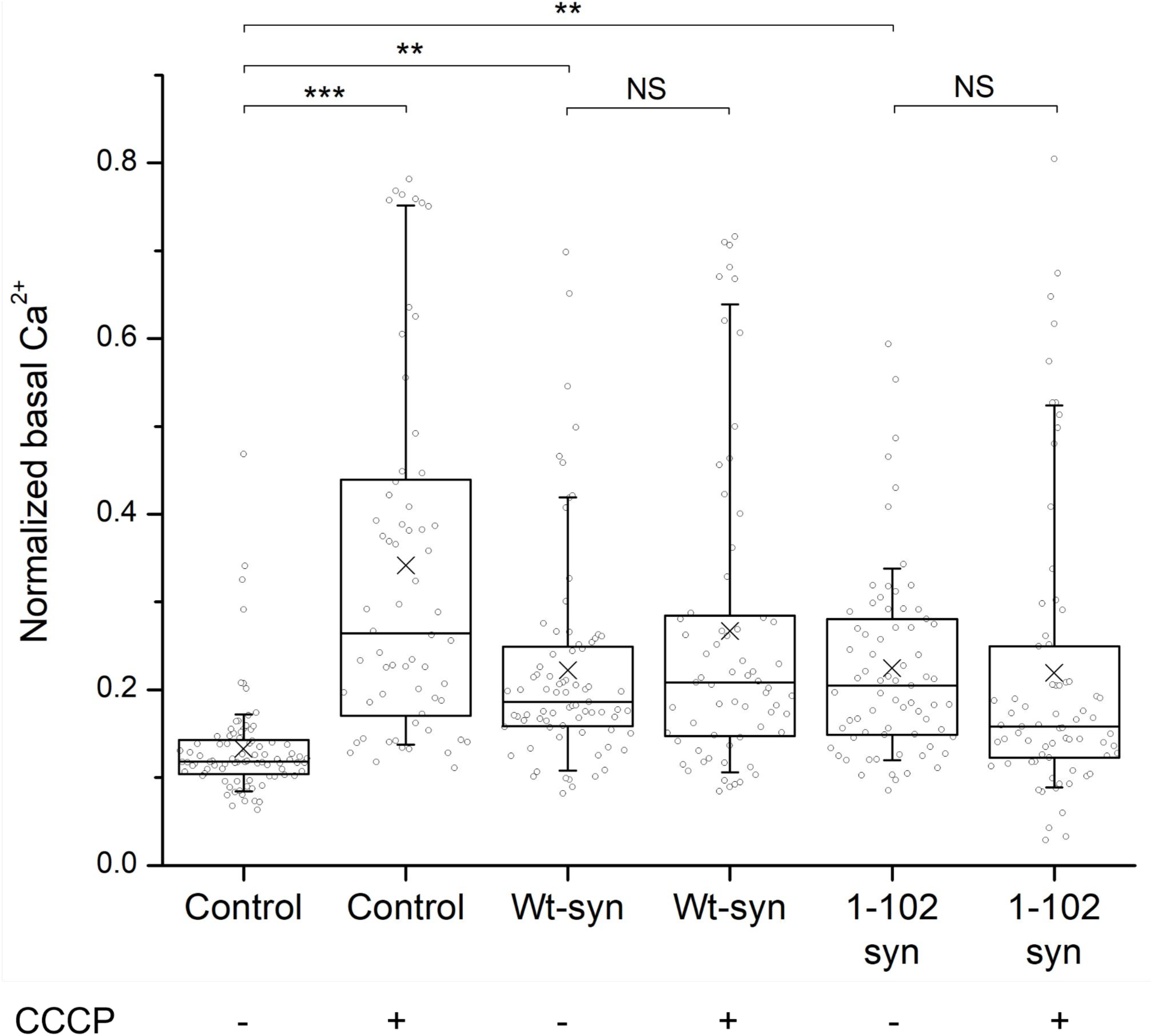
Expression of Wt-syn and 1-102-syn enhances basal level of mitochondrial Ca^2+^ with or without CCCP treatment. RBL cells were co-transfected with mito-GCaMP6f and one of pcDNA (control), Wt-syn or 1-102-syn. Samples were treated or not with 10μM CCCP for 30 min, then washed with RBL media and incubated in media for 3 hours (recovery), followed by imaging using confocal microscopy. Mito-GCaMP6f fluorescence was monitored in confocal movies initially and after addition of high-dose ionomycin (5 μM) to achieve maximal fluorescence. Each sample comprising n∼100 co-transfected cells are from three independent experiments. The box plots show 25th-75th percentile of the data; midline shows median, and X shows average; ** represents P- values <0.01, *** represents P-values <0.001. NS represents not significant (P-values > 0.05).

<u>Movie S1. RBL cells transfected with Wt-syn exhibit high level of mitochondrial Ca^2+^ uptake when stimulated with sub-optimal concentration of antigen.</u> Movie corresponds to trace shown in Figure 2b of main text; image field includes about 25 cells. RBL cells were co-transfected with plasmids for mito-GCaMP6f and Wt-syn and imaged by confocal microscopy. Mitochondrial Ca^2+^ uptake was monitored mito-GCaMP6f fluorescence before (starting at -16 sec) and after stimulation by Ag (DNP-BSA, 1 ng/ml, starting at 0 sec), followed by addition of high-dose ionomycin (5μM, starting at 370 sec).

<u>Movie S2. RBL cells transfected with empty vector exhibit little mitochondrial Ca^2+^ uptake when stimulated with sub-optimal concentration of antigen.</u> Movie corresponds to trace shown in Figure 2c of main text; image field includes about 25 cells. RBL cells were co-transfected with plasmids for mito-GCaMP6f and pcDNA (empty vector control) and imaged by confocal microscopy. Mitochondrial Ca^2+^ uptake was monitored mito-GCaMP6f fluorescence before (starting at -16 sec) and after stimulation by Ag (DNP-BSA, 1 ng/ml, starting at 0 sec), followed by addition of high-dose ionomycin (5μM, starting at 370 sec).

